# Structural features of *E. coli* Stx bacteriophage phi24B revealed with cryo-electron microscopy

**DOI:** 10.64898/2026.04.10.717836

**Authors:** Matvey A. Bubenchikov, Alexander S. Kuznetsov, Ivan O. Butenko, Darya S. Matyushkina, Olga S. Sokolova, Andrey V. Letarov, Andrey V. Moiseenko

**Author notes:** **Author Contributions:** M.A.B. performed cryo-EM data processing, structural modeling and manuscript writing. A.S.K. carried out phage purification, virological studies and manuscript writing. D.S.M. and I.O.B. performed proteomics experiments. O.S.S. acquired cryo-EM data. A.V.L. performed conceptualization, project administration, data analysis and manuscript writing. A.V.M. performed funding acquisition, project administration, manuscript writing and contributed to cryo-EM data processing. **Competing Interest Statement:** No competing interest.

## Abstract

Shiga toxin-converting bacteriophages play a critical role in the emergence and virulence of pathogenic Escherichia coli strains. Despite their significance, detailed structural information on these phages remains scarce. Here we present a high-resolution cryo-electron microscopy and proteomic analysis of the phi24B bacteriophage, revealing an icosahedral capsid with T=9 symmetry, decorated by a processed esterase protein (gp84) and stabilized by cementing proteins. The tail assembly comprises a dodecameric portal, two rings of adapter proteins sharing a common fold, a hexameric nozzle, six lateral tail fibers, and a flexible central needle fiber. The binding sites of the fibers are described. Comparative analysis indicates conservation of the tail structure with related podoviruses but very different peripheral features.

## Introduction

Viruses of bacteria – bacteriophages (phages) – not only contribute to daily mortality of bacterial cells in various natural microbial communities (DiPietro et al. 2023; Heinrichs et al. 2020) but also strongly impact the physiology and evolution of their hosts (Liang et al. 2025; A. V. Letarov and Letarova 2023). Temperate phages, able not only to grow in lytic cycle, but also to integrate bacterial genomes and to exist in the form of prophages in lysogenic cells, are long known to contribute to the pathogenicity of many species of bacteria (Howard-Varona et al. 2017). The most understood mechanism for this is lysogenic conversion when some genes carried on a prophage are expressed in the lysogenic cell providing it with some features that improve the host fitness under certain conditions. Pathogenicity factors are among the most known converting genes (Howard-Varona et al. 2017; Cieślik et al. 2021), and the key factor that determines the virulence of the pathogenic bacteria is usually toxin production. Prophages are responsible for synthesis of many bacterial toxins, including diphtheria toxin (Freeman and Morse 1952; Freeman 1951), *Staphylococcus aureus* hemolytic exotoxin (Y. Li et al. 2025), one of the *Clostridium botulinum* toxins (Barksdale and Arden 1974), cholera toxin (Waldor and Mekalanos 1996), *Pseudomonas aeruginosa* cytotoxin CTX (Nakayama et al. 1999) and some other toxins. In most of these cases the toxins are expressed in the lysogenic cells and secreted by bacterial secretion systems so the lysogenic bacteria become bona fide toxigenic cells.

The production of Shiga-like toxins (Stx) by the Shiga-toxigenic *Escherichia coli* (STEC) strains is also dependent on the presence of Stx-converting prophages. However, the genes of the Shiga-toxin subunits StxA and StxB are only transcribed during the lytic development of the phage, and the toxin is released by the cell lysis alongside the phage progeny (Rodríguez-Rubio et al. 2021). Therefore, the individual lysogenic cells are not toxigenic but the lysogenic population produces a certain amount of the toxin due to spontaneous prophage induction of a fraction of the cells. The toxin enters the bloodstream and damages epithelium of the gut and kidney blood vessels. The latter may cause so-called hemato-uremic syndrome (HUS), a life-threatening condition that often results in long-term disability in many surviving patients (Nakao and Takeda 2000). The development of HUS in particular patients depends on the toxin subtype and the level of the toxin load (Nakao and Takeda 2000). Notably, the STEC infections are normally self-limiting and the pathogen gets spontaneously eradicated in about two weeks. Therefore, the treatment of STEC infection mostly relies on supportive therapy while the administration of the antibacterial drugs may enhance the HUS incidence since the stress caused by them in the pathogen’s cells may increase the induction level and toxin production (Sevilla et al. 2025).

The lysogenization of *E. coli* strains featuring moderate pathogenic potential by Stx-converting phages may turn them into a serious pathogen causing dangerous foodborne infections. Although the most known STEC serotype is O157:H7 the other non-O157 STEC serotypes are gaining epidemiological significance. For example, an O104:H4 strain has caused in 2011 a major outbreak in Germany with 3842 cases in 22.3% of which HUS development was reported (Beutin and Martin 2012). In addition to the most known O157:H7 and O104:H4 serotypes about 400 of different STEC serotypes were associated with infections in humans, among them so-called “Big six” (O26, O45, O103; O111, O121 and O145) have major epidemiological significance (Mellor et al. 2016). The heteropathogenic enterohemorrhagic – uropathogenic E. coli (EHEC-UPEC) strains were also described (Toval et al. 2014). Therefore, the ability of the Stx-phages released by STEC pathogens to infect and lysogenize environmental *E. coli* strains is believed to be a major driver of the emergence of new STEC strains. The lytic multiplication of the Stx phage on the commensal *E. coli* within a patient may also contribute to the toxin load and influence the outcome of the particular case, the implementation of this process was demonstrated in a mouse model (X. Zhang et al. 2000). These considerations highlight the importance not only of the Stx-prophage physiology (for example, the factors influencing the induction levels) but also the structure of the virions and infection mechanisms of Stx phages.

From the genetic organization perspective, almost all the Stx-phages belong to the lambdoid group, sharing overall genome organization and core of the lysogeny control mechanism with bacteriophage λ (Maite Muniesa et al. 2004; Rodríguez-Rubio et al. 2021). At the same time, their virions span all major morphological classes of tailed bacteriophages—including podoviruses, siphoviruses, and myoviruses (M. Muniesa et al. 2004), highlighting the modular nature of lambdoid phage evolution (Botstein 1980; Canchaya et al. 2003). More than half of the known Stx phages are related to the bacteriophage vb_EcoP_24B (also known as phi24B).

The bacteriophage phi24B belongs to the species *Traversvirus* tv24B within the genus *Traversvirus* in the subfamily *Sepvirinae* in the class *Caudoviricetes*. It is a lambdoid podovirus with a genome of 57,677 bp which is larger than that of most lambdoid phages (for example, 48,502 bp in phage λ). In addition to the StxAB genes located downstream of the late gene antiterminator Q, phi24B encodes the gp84 protein within its late gene cluster. This protein contains a large domain closely related to the *Escherichia coli* deacetylase NanS, which removes acetyl groups from mucin O-glycans (Franke et al. 2020). It has been proposed that this gene enables lysogens to utilize the mucin as a carbon source (Rodríguez-Rubio et al. 2021). In addition to lysogenic conversion, the carriage of the phi24B prophage has been reported to alter the metabolism of the lysogenic cells through the unknown mechanism, enhancing their survival in the gastrointestinal tract (Holt et al. 2017) and potentially facilitating STEC transmission.

The host range of phi24B defined as the repertory of *E. coli* strains supporting the phage plaques formation is limited and includes mainly rough strains such as laboratory *E. coli* K12 derivatives (James et al. 2001; Golomidova et al. 2021). At the same time, the lysogenization range is apparently broader than was shown by the usage of genetically marked phi24B:Cat strain of the phage forming antibiotic-resistant lysogens out of many wild *E. coli* strains that do not support plaque formation (James et al. 2001). Such a broad infection range may be explained by the fact that phi24B uses a highly conserved outer membrane protein BamA (previously YaeT) as the receptor (Smith et al. 2007). However, most of the wild *E. coli* strains produce O antigens efficiently screening the cell surface proteins from direct interaction with phage RBPs and bacteriophages need to acquire specific proteins to penetrate the O antigen barrier (A. V. Letarov 2023). The analysis of the lysogens formed by phi24B:Cat under the laboratory conditions out of environmental O-antigen-protected E. coli strains revealed that all the lysogens checked were depleted of the O-polysaccharide (OPS) or featured decreased OPS synthesis (Golomidova et al. 2021) but the presence of the prophage did not interfere per se with the OPS production. Therefore, it was concluded that phi24B lysogenization as well as its lytic infection is efficiently blocked by the O-antigen barrier. At the same time many natural STEC strains carrying prophages closely related to phi24B produce various types of the O-antigens posing the question on the natural mechanism(s) involved in the transmission of these phages.

The structure of phi24B virion and its structural proteome were not yet characterized in detail and it is not known which protein is responsible for the interaction with the BamA receptor and if some other receptor recognition proteins are displayed on the virion and may be involved into the interaction with some O-antigens or other molecules (A. V. Letarov 2023) that could facilitate OPS layer penetration. The lack of structural data for phi24B was probably due to the problematic procedure of culturing and purification of this virus hindered production of high concentration stocks suitable for cryo-electron microscopy (cryo-EM) investigation. Previously we managed to produce highly purified samples of phi24B by induction of the lysogen obtained from the strain E. coli 4sR with subsequent sucrose gradient purification (Kuznetsov et al. 2024). Here we present the high-resolution cryo-electron microscopy study of structural organization of phi24B virions.

## Results

### Virion protein composition revealed by proteomics

To characterize the structural proteome of bacteriophage phi24B we run the protein gel (Fig. 1A) of the sample used for cryo-EM data collection. Protein bands were excised (Fig. S1) and identified using trypsin peptidomics. The total phage suspension was also subjected to trypsinolysis and peptide profiling in a batch proteomic experiment.

**Figure 1.**
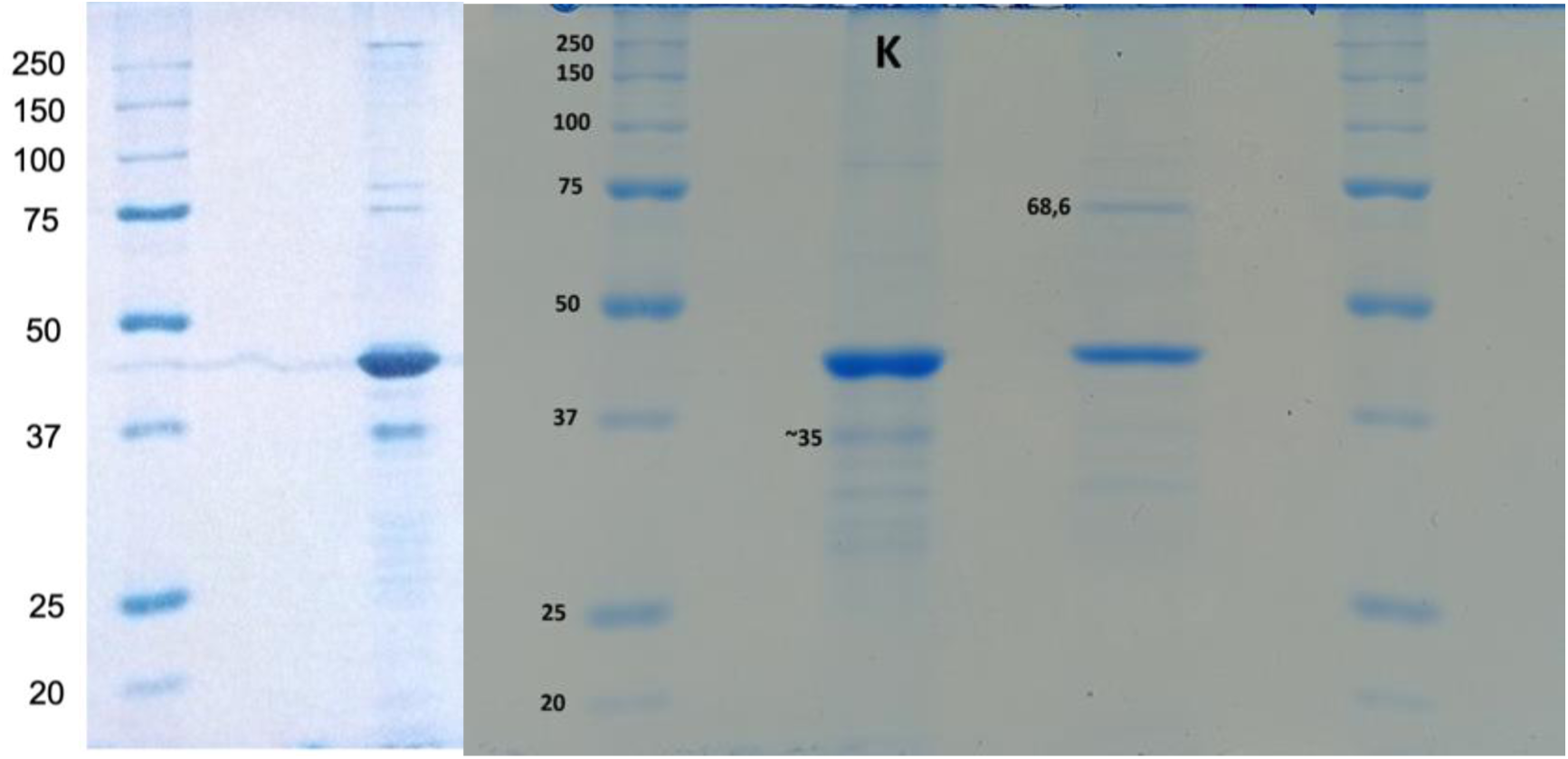
Protein profile of phage phi24B samples. A. Protein gel used for the band identification by MS-proteomics. Left lane, protein molecular weights marker; right lane, phi24B phage proteins. B. Comparison of the same sample shown in panel A (left lane) with a freshly purified phage preparation (right lane).

The total phage suspension was also subjected to trypsinolysis and peptide profiling in a batch proteomic experiment. Fifteen phage-encoded candidate structural proteins were identified (Table 1), including all the proteins resolved in the cryo-EM experiment (see Results section below). Four additional phage encoded proteins, although detected at signal levels comparable to low abundance structural proteins, were classified to be contaminating non-structural protein based on their functional annotation (a transcription regulation factors, serum-resistance proteins Bor and Lom and a homolog of a bacteriocin).

**Table 1.** Estimated and structure-derived abundances of φ24B proteins. Abundances are normalized to the portal protein and compared with values inferred from the structural model.

| Protein length, a.a. | Protein ID and function | Abbreviation | Estimated abundance, copies per virion | Structure-based abundance, copies per virion |
| --- | --- | --- | --- | --- |
| 2811 | gp47 ejection protein | EjP | 23 | ND |
| 724 | gp72 Portal | PrP | ND | 12 |
| 665 | gp84 carbohydrate esterase decorating capsid protein | DCP | 31 | ND |
| 649 | gp61 tail fiber | TF | 22 | 18 |
| 579 | gp57 nozzle protein | NZP | 6 | 6 |
| 460 | gp56 tail needle protein | TNP | 6 | 3 |
| 421 | gp48 hypothetical protein | – | 37 | ND |
| 409 | gp69 main capsid protein | – | 352 | 535 |
| 342 | gp70 hypothetical protein | – | 9 | ND |
| 260 | gp51 hypothetical protein | – | 27 | ND |
| 234 | gp64 tail tube (adaptor 1) | ADP1 | 11 | 12 |
| 245 | gp62 tail tube (adaptor 2)1 | ADP2 | 2 | 12 |
| 161 | gp50 hypothetical protein | – | 44 | ND |
| 202 | gp54 putative receptor binding | RBP | 2 | ND |
| 150 | gp67 capsid cementing protein | CCP | 290 | 525 |
| 75 | gp55 putative receptor binding protein | - | Not found | Not found |

Using peptide intensity measurements from the batch proteomics experiment, together with copy numbers inferred from the structural model for selected reference proteins, we estimated the abundance of each detected protein. Abundances normalized to the portal protein (assumed to be present at 12 copies per virion) are shown in Table 1. Comparison with structure-derived stoichiometries indicates that, for most proteins, the estimated abundances deviate by a factor of 1–3 (Table S1).

Among the detected proteins, gp47 is notable for its large size 2811 amino acids. Conserved domain analysis identified LPD5, LPD1, and LPD38 domains characteristic of phage large polyvalent proteins, as well as a Muf-C metallopeptidase domain. Despite its abundance, gp47 was not observed in cryo-EM or negative-stain images and is absent from the reconstructed virion model (see below). We therefore propose that gp47 is packaged within the capsid and functions as an ejection protein, forming or contributing to a transperiplasmic conduit that extends the bacteriophage tail during the host cell infection. The metallopeptidase domain likely facilitates degradation of the peptidoglycan layer, enabling this conduit to reach the plasma membrane of the cell.

Another protein unexpectedly detected in the virion is gp84. This protein contains a highly conserved esterase domain related to the *E. coli* NanS enzyme, which is required for the mucin utilization as a carbon source for bacterial growth (Franke et al. 2020). It has been proposed that gp84 may act as a conversion protein improving the lysogen fitness (Rodríguez-Rubio et al. 2021).

One of the most intense protein bands in the phage protein profile was identified as gp84; however, its apparent molecular weight of about 35KDa was substantially lower than the predicted 68.4 KDa for the full-length protein. Notably, the phage preparation used for proteomic analysis had been stored at 4 °C for six months, although it retained high infectivity. This observation is consistent with previously reported autoproteolysis of purified gp84, resulting in cleavage of the C-terminal jelly-roll domain (Franke et al. 2020) (Fig. 2). To verify this, we analyzed a freshly purified phage preparation alongside the stored sample (Fig. 1B). In the fresh sample, the ∼35 kDa band was absent, and instead a new band at ∼70 kDa was observed. The MS based peptidomics confirmed that this larger band contains gp84 protein. Analysis of the peptides detection confirmed that the peptides mapping to the C-terminal domain were only detected in the 70KDa band once the middle deacetylase domain was detected in both 35KDa and 70KDa bands. These results indicate that gp84 is a structural protein undergoing proteolytical processing after the release of the phage from the cell.

**Figure 2:**
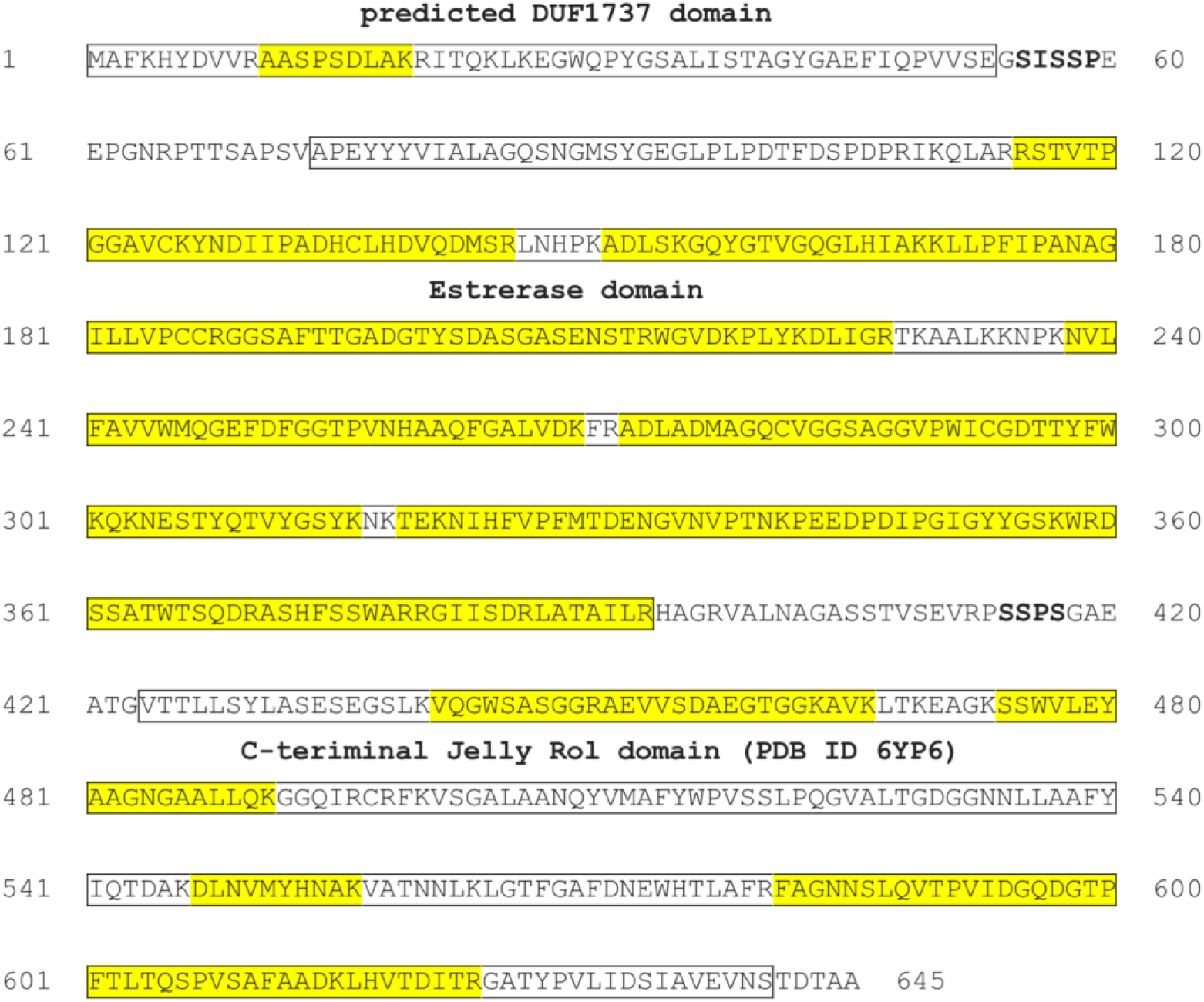
Sequence and domain structure of the gp84 protein. Peptides identified by LC-MS analysis are highlighted in yellow. Putative motifs involved in autoproteolytic processing are indicated in bold.

Given that the polysaccharide deacetylase domain would require exposure on the virion surface to access its substrate, we suggested that gp84 may be a capsid decorating protein. Noteworthy, gp84 band is one of the most intensive bands in the phage protein profile after the MCP gp69 band (Fig. 1), indicating high copy number of this protein in the virion (potentially exceeding the ∼31 copies estimated from peptide counts; Table 1).

We therefore searched for corresponding density in the cryo-EM maps. This analysis revealed a well-resolved density corresponding to a hexamer of the small N-terminal domain of gp84, located within the central cavities of selected capsid hexons (see below). Together these observations identify gp84 as a decorating capsid protein. Notably, the density corresponding to the N-terminal domain terminates abruptly after residue Gly54. Even if the central NanS-related domain were disordered and averaged out in our reconstruction, the hexamer of 30 KDa domain (total molecular weight of 180 KDa) would be expected to produce detectable density. The absence of such density suggests that gp84 undergoes additional proteolytic cleavage not only between the C-terminal and central domains esterase but also between the central domain and N-terminal capsid-binding domain (previously annotated and a domain of unknown function DUF1737). We manually inspected the protein sequence of the interdomain linkers in gp84 (Fig. 2) and found only SSP-containing motifs to be similar. The left SISSP sequence is closely adjacent to the last visible Gly54 residue.

### The structure of phi24B virion solved with cryoEM

#### Overall virion composition

We performed cryo-electron microscopy analysis of phi24B virions. The dataset comprised projections of approximately 88,000 individual particles. Image inspection revealed that ∼52,000 particles corresponded to intact virions displaying morphology typical of podoviruses, whereas the remaining ∼36,000 particles exhibited capsids lacking genomic DNA (Fig. 3A). We suggest that residual host cell debris present during the early stages of purification may have triggered DNA ejection through interactions between phages and cellular antigens. We therefore refer to these virions as the ejected state.

**Figure 3:**
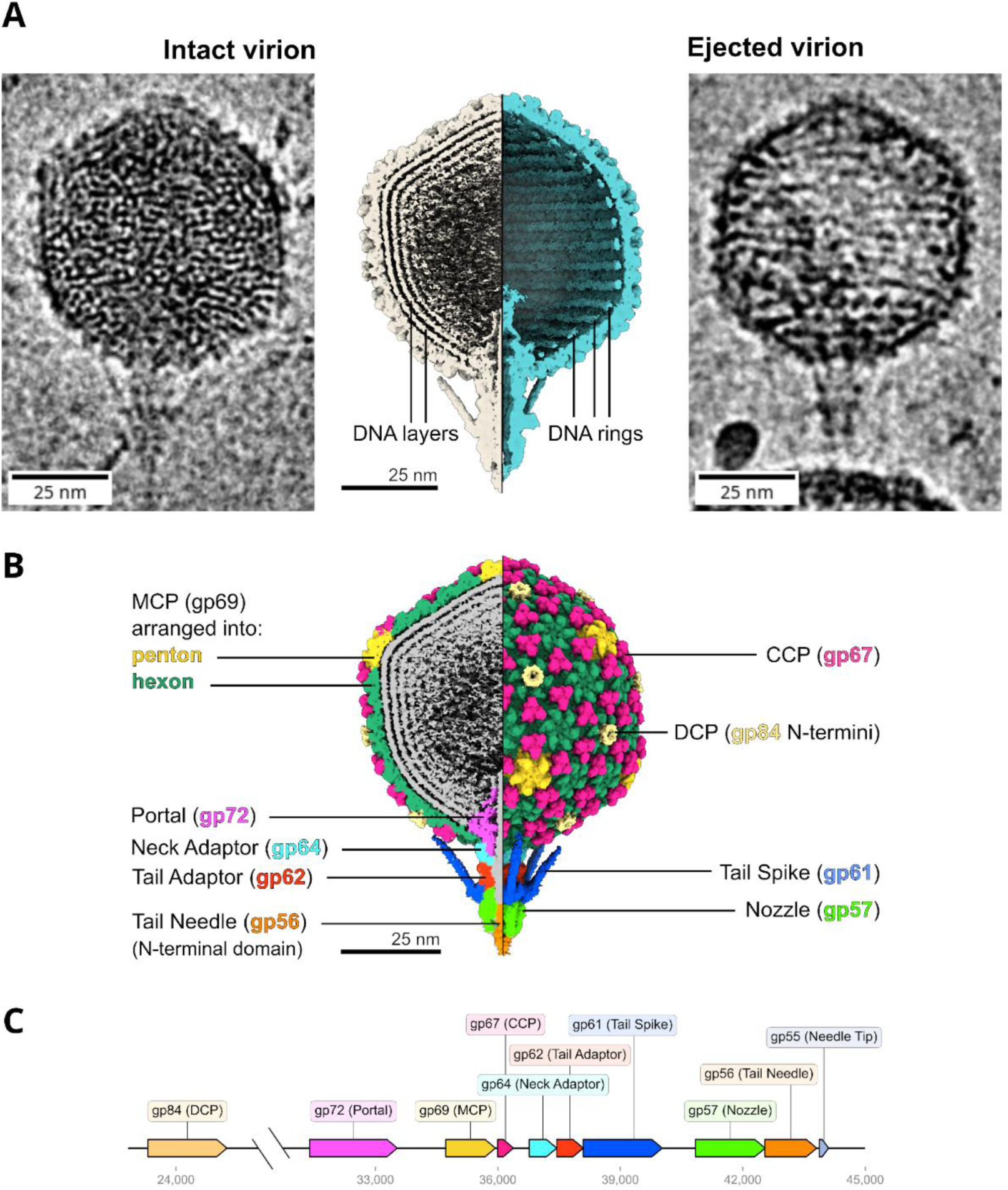
Overall composition of the phi24B virion. A - Representative cryo-EM images and side-by-side sections through the three-dimensional reconstructions of intact (left) and ejected (right) phi24B virion. B - Asymmetric cryo-EM reconstruction of the intact phi24B virion colored according to protein composition. C – Genomic organization of phi24B structural proteins. Numbers indicate genome coordinates in base pairs.

Using single-particle analysis, we reconstructed the three-dimensional structure of the virions and built molecular models for most proteins forming the capsid shell, portal complex, and adsorption apparatus, as well as partial models of the fiber proteins (Fig. 3B,C). We have performed the same analysis for both intact and ejected virions. The asymmetric three-dimensional reconstruction of the phi24B is shown in Fig. 3A,B.

The virion contains an isometric icosahedral capsid approximately 74 nm in diameter (vertex to vertex) with a triangulation number of T = 9, a relatively rare symmetry among podoviruses that has previously been observed in *Pseudomonas* phages Moo19, B2, distantly related to the *E. coli* phage N4 (Subramanian et al. 2024), and in *Ralstonia* phage GP4 (Zheng et al. 2022, 2023). The phi24B tail is short (∼22 nm) and carries six lateral fibers and a single central fiber. The sections through three-dimensional density maps of intact phi24B virions reveal four distinct concentric layers of tightly packed DNA within the capsid (Fig. 3A, B). In contrast, these concentric layers of DNA are missing from the ejected state, but a single layer of DNA remains within the capsid after genome release, likely retained through interactions with the inner surface of the capsid shell (Fig. 3A).

In intact virions, the portal channel contains a continuous cylindrical density extending from the capsid interior, which we assign to genomic DNA by analogy with previous cryo-EM studies of bacteriophages (Bayfield et al. 2023; Chen et al. 2021; Molineux and Panja 2013; Zheng et al. 2022). The density protruding from the distal end of the tail corresponds to the central fiber protein gp56, as described below. Neither the cylindrical DNA density within the portal channel nor the density corresponding to the central fiber is observed in the ejected state.

Asymmetric reconstruction of the phi24B virion indicates that the symmetry-mismatched interface between the C5 capsid vertex and the C12 portal complex has a stable conformation. Analysis of conformational heterogeneity revealed two distinct orientations of the tail assembly relative to the capsid (Fig. S3B). In ejected virions, the tail is rotated by approximately 6° about its longitudinal axis compared with intact particles, consistent with previous observations for phage Sf6 (F. Li et al. 2022). In addition, intact volumes exhibit a slight tilt of the tail body of ∼2° relative to the central axis between the two conformational states. We also obtained four focused cryo-EM reconstructions of the asymmetric neck regions in which both portal and capsid symmetries are resolved simultaneously (Fig. 5D). The resolution about 4.3 Å allows reliable tracing of individual protein chains and detection of rotational differences of ∼6° in the tail assembly. A tilt of ∼2° of the portal relative to the capsid axis is observed in both states. However, consistent with the whole-virion reconstructions, the tail body is tilted only in intact particles, whereas in ejected virions it aligns with the central axis (Fig. S3B). This change in tail alignment may be associated with the presence or absence of packaged DNA and internal tail contents, which could act as an internal “core” constraining the tail orientation relative to the portal. The asymmetric neck reconstructions further reveal the symmetry-mismatched interactions between the dodecameric portal and the pentameric capsid vertex in tailed phages. However, the achieved resolution is insufficient for detailed atomic modeling or unambiguous identification of side-chain interactions.

Inspection of raw micrographs revealed an unusual orientation bias of phi24B virions relative to the vitrified ice layer. Based on the geometry of the virion, particles embedded in the thin aqueous layer spanning the holes of the carbon support would be expected to adopt orientations with the tail aligned parallel to the ice plane. However, consistent with the 2D classification of the phi24B virions (Supplementary Fig. S3A), approximately 66% of intact particles and 75% of ejected virions were oriented with their tails pointing toward the ice surface. This orientation likely reflects interactions between components of the adsorption apparatus and the air–water interface prior to sample vitrification.

#### Structure of the capsid proteins

To determine the structure of the capsid proteins, we performed three-dimensional reconstruction of the capsid shell imposing icosahedral symmetry. The resulting maps reached resolutions of 3.8 Å for intact virions and 4.0 Å for ejected virions (Fig. S5). We suggest that the relatively large capsid diameter (∼74 nm) may lead to minor deviations from perfect icosahedral symmetry caused by embedding in the vitrified ice; thereby limiting the resolution of the symmetrized reconstructions. To address this, we carried out local reconstructions of smaller regions of the capsid shell, yielding three overlapping density maps that together cover the asymmetric unit (Fig. S5). These local reconstructions reached resolutions of 2.5–2.7 Å, which was sufficient for reliable atomic model building.

The capsid shell is composed of three proteins: the major capsid protein gp69 (MCP), the cementing capsid protein gp67 (CCP), and the N-terminal fragment of the processed gp84 protein, which functions as a decorating capsid protein (DCP) (Fig. 4A). Accordingly, each asymmetric unit of the icosahedral capsid contains nine MCP, nine CCP, and two DCP subunits.

**Figure 4:**
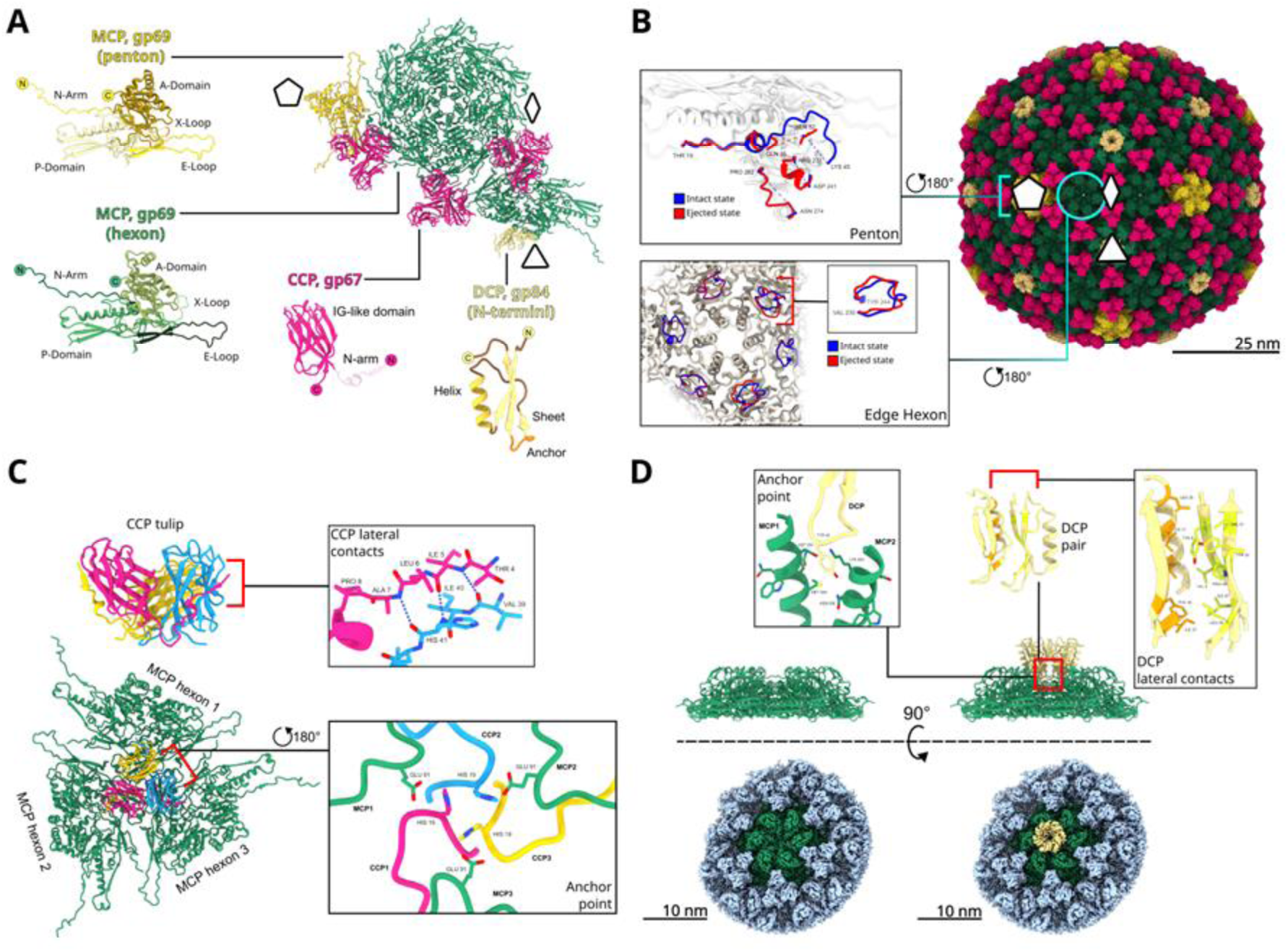
Capsid reconstruction. A – One asymmetric unit and individual chains of capsid proteins colored by protein chains. Pentagon, Triangle and rhombus indicate the positions of the symmetry axes. B - Molecular model of icosahedral capsid and focused views from the inside of capsid interior on superimposed models in both Ejected and Intact states. C - Overview of CCP (gp67). CCP trimer with contact between N-arm and Ig-like domain of two monomers, three-capsomer contact points are shown. D - Structural features of DCP (gp84 N-termini). Atomic models and densities of Face hexons with and without DCP hexamer, contacts between DCP monomer and pair of axial MCP helices and between two DCPs are shown.

The major capsid protein adopts an HK97-like fold and assembles into capsomers, a structural motif that predominates among tailed bacteriophages. Each MCP subunit comprises an N-terminal arm, an E-loop, a P-domain, and an A-domain (Fig. 4A). The N-terminal arm forms an interaction between neighboring MCP subunits within a capsomer, which is typical for HK97-like capsid architectures (Wikoff et al. 2000). In contrast to many HK97-like phages (Wikoff et al. 2000; Šiborová et al. 2022), the N-terminal arm of phi24B MCP shows no evidence of proteolytic cleavage and is resolved in the density map starting from residue Thr2. A similar feature is also discovered in phages GP4, Pam1, T7, Moo19, B2, Sf6 (J.-T. Zhang et al. 2022; Guo et al. 2014; F. Li et al. 2022; Subramanian et al. 2024).

The phi24B MCP lacks inter-subunit isopeptide bonds between E-loops of neighboring capsomers. In many HK97-like phages (Zhou and Chiou 2015) such covalent linkages form a “chainmail” network that enhances capsid stability. In phi24B there are only non-covalent, electrostatic interfaces.

The closest structural homolog of phi24B is *Ralstonia solanacearum* podovirus GP4. The structures of the phi24B MCP gp69 and the GP4 MCP gp2 are highly similar, consistent with their sequence identity of 32% and E-value of 3.7e-39 based on BLASTp search with BLOSUM62 matrix (Fig. S2). In phi24B, the P-domain contains an additional loop spanning residues Ala37-Pro56 that is absent in GP4. Although this loop extends toward the capsid interior, its density is not resolved in the reconstruction, indicating conformational flexibility. This loop contains four lysine residues (Lys39, Lys45, Lys46 and Lys69), suggesting a potential role in interactions with the DNA packaged in capsid.

In the penton form of the MCP, regions Gly35-Ser52, Arg231-Asp241, and Asn274-Pro282 are not resolved in the intact state, but are resolved in the ejected state (Fig. 4B and S3D). All three regions extend from the inner surface of the capsid shell toward the capsid interior. We propose that these flexible loops participate in interactions with the packaged DNA, that are lost after the DNA release. Several additional regions of the MCP are only partially resolved in the density maps. All of these segments extend toward the capsid interior. In particular, residues Gly348-Gly353 lack interpretable density in both intact and ejected states across all MCP subunits.

Another difference between intact and ejected states in MCP structure is found in two MCPs participating in the edge hexon. The two that are on opposite sides of the inflection edge line and near to the penton have notable differences in the loops Val230-Glu244 that protrude to the capsid interior. In intact virions these loops are packed as a one-step alpha helix, like it is found in other hexon MCPS. At the same time, in ejected virions the loops are not behaving as similar loops in hexon, the structure is planar (Fig. 4B and S3D).

The capsid cementing protein gp67 assembles into trimers, each binding to the outer surface of the capsid at sites where three neighboring capsomers meet (Fig. 4C). Each CCP subunit comprises an N-terminal arm followed by a 4+4 immunoglobulin-like domain. The N-terminal arm forms a β-sheet with a neighboring CCP subunit, creating a “hugging” interface that stabilizes the trimer into a rigid, tulip-like structure. The CCP trimer is anchored to the capsid shell by three His19 residues directed toward the threefold interface formed by neighboring MCP capsomers. A similar fold, anchoring mode, and “hugging” interface have been reported in bacteriophage PAM1 (sequence identity of 21%) (J.-T. Zhang et al. 2022). Although the cementing protein of GP4 phage also adopts an Ig-like fold and anchors at the three-capsomer junction, it uses Asp residues for anchoring and lacks the “hugging” inter-subunit interface.

At the center of each icosahedral capsid face—that is, at the central hexon formed by MCP subunits—we observe an attached hexamer of a small protein, which we assign to the N-terminal domain of the gp84 esterase and refer to as a decorating capsid protein (DCP). This domain corresponds to the previously described domain of unknown function DUF1737, homologues of which are present in several bacteriophages (Franke et al. 2020).

The structure of gp84 esterase was previously reported in (Franke et al. 2020) and comprises three domains: a large catalytic and jelly roll domains – both contributing to enzymatic activity, and the small DUF1737 domain, which we identified here as DCP. The structure of the observed N-terminal domain consists of an alpha helix and three antiparallel beta strands connected by two loops: one between the helix and sheet and another acting as an anchor with Tyr42 residue that plays as a “tooth” stuck in a cavity formed by two MCP helices of central pore. (Fig. 4D).

The resolution of our density map enabled unambiguous modeling of residues Ala2-Ser57 of gp84, whereas the remaining portion of the protein is not resolved. The catalytic and jelly-roll domains together account for a substantial molecular mass (∼63 kDa), which would correspond to ∼380 kDa in a hexameric assembly. Even if this region were flexible, it would be expected to generate detectable unstructured density in the cryo-EM reconstruction; however, no such density is observed. We therefore propose that gp84 undergoes proteolytic cleavage after residue 57, leaving only the N-terminal DUF1737 domain associated with the capsid.

Notably, according to the Pfam/InterPro databases (Blum et al. 2025; Paysan-Lafosse et al. 2025), the DUF1737 domain (PF08410) is found in approximately 3000 proteins (Fig. S9), 2528 of them are single-domain proteins, 481 proteins are supposed to be estarases, because they consist of DUF1737 and SASA domains, and others have beta-lactamase, isochorismatase or other domains. There are around 2000 viral and bacterial species genome sequences with the presence of DUF1737 domain in encoded proteins, that are associated, among others, with prophage genes.

The gp84 hexamer binds to the pore at the center of a capsomer formed by six adjacent MCP subunits (Fig. 4D). Notably, gp84 is found exclusively on hexons located at the centers of icosahedral faces and is absent from other hexons. We suggest that this selectivity is caused by local capsid shell curvature: central hexons are relatively flat, whereas hexons near edges and vertices are slightly bent and may prevent gp84 binding. Three-dimensional classification further revealed partial occupancy of gp84 across the capsid surface. Approximately 80% of capsid faces in intact virions and ∼50% in ejected virions are decorated with gp84 hexamers, whereas the remaining faces lack gp84 entirely. No states corresponding to alternative gp84 oligomeric states were detected.

Not all bacteriophages possess decorating proteins at the centers of capsid hexons. The closely related phage GP4, for example, encodes only a single type of decorator protein. However, a structurally similar decorating protein has been described in phage Bas63 (Hodgkinson-Bean et al. 2025). In Bas63, a protein occupies the center of each hexon and forms “teeth”-like interactions with cavities of the MCP surrounding the central pore. The monomer adopts a fold resembling the globular domain of the phi24B DCP, comprising a short α-helix and an antiparallel β-sheet. A difference lies in oligomeric organization: whereas phi24B assembles its DCP into hexameric rings, Bas63 features a single asymmetric monomer at the center of each capsid face.

#### The portal protein and tail structure

We have built the cryoEM reconstructions for phi24B portal and tail regions of the phi24B virion in both intact and ejected states. Multiple reconstructions focused on different parts of the tail yielded the density maps with resolutions in range 2.7 to 5.1 Å (Fig. S5). We have built the molecular models for all of the proteins that constitute the tail assembly (Fig. 5A). The resolution values of obtained densities were enough to collect insights on sidechains orientations in contacts of distinct rings. The interfaces are elaborated, including salt bridges, hydrogen bonds and hydrophobic interactions. The contacts are highly individual for each pair of rings and form highly specific cavities and bumps (Fig. S7).

**Figure 5.**
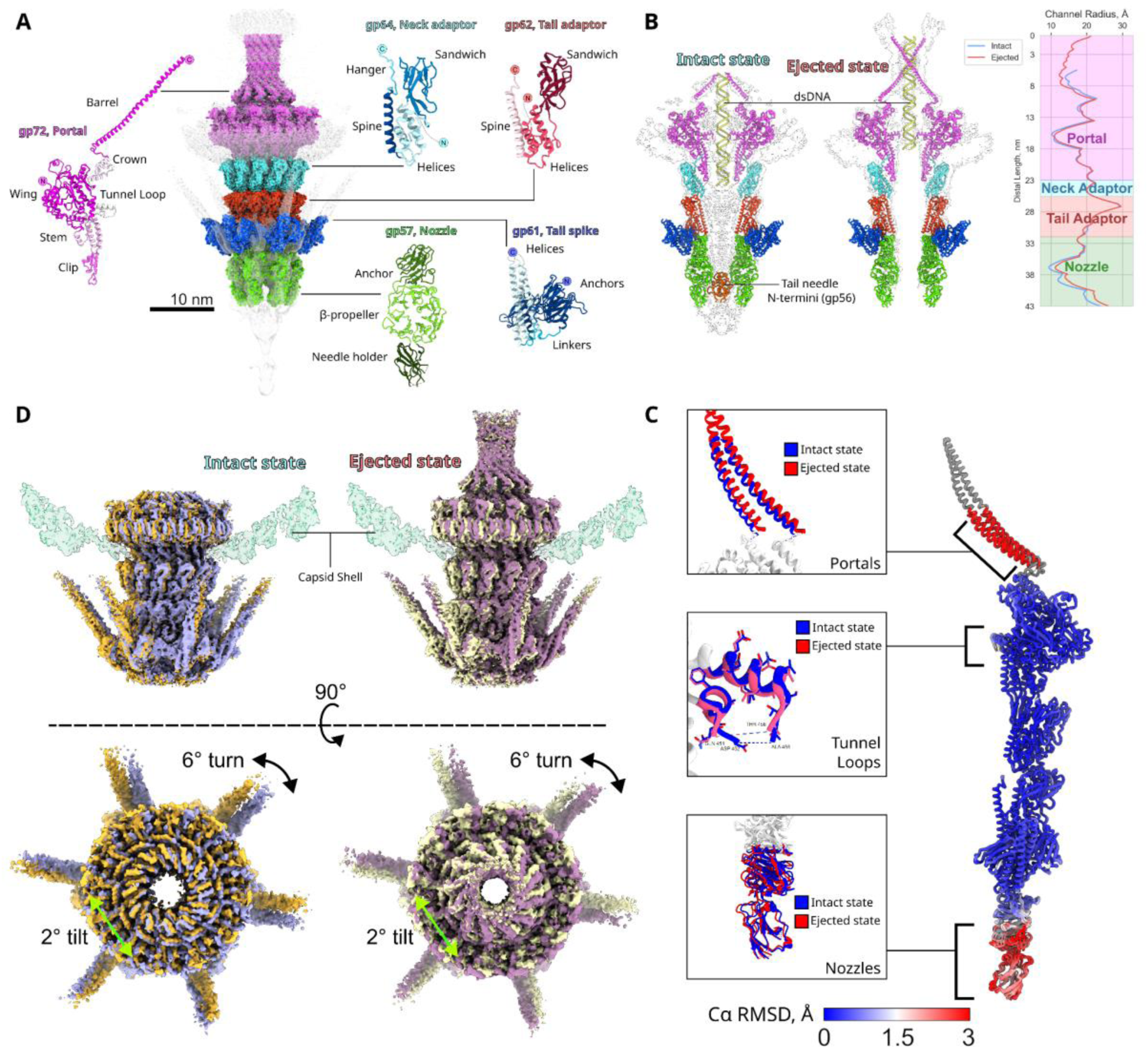
phi24B tail composition and rearrangement upon DNA release. A – Molecular model of the tail assembly shown as surface and colored by proteins. Individual monomers are shown as ribbons and colored by domain. B – Central cross-section of tail molecular model in intact and ejected states with HOLE (Smart et al. 1996) analysis. The density maps are displayed semi-transparent. C - Cα RMSD between asymmetric units of the intact and ejected states highlighting the regions of differences. D - side and top views on Capsid-Portal asymmetric interface with superimposition of conformers in Intact and Ejected states

The overall architecture of the phi24B tail, proceeding from the capsid toward the distal tip, is as follows. A dodecameric portal protein, gp72, is incorporated into the capsid shell by replacing one of the MCP pentons. Two concentric dodecameric rings of adaptor proteins, gp64 and gp62, are attached to the portal complex, followed by the hexameric nozzle protein, gp57, which completes the tail structure. Six lateral fibers, each formed by a gp61 trimer, are attached to the point of contact of nozzle and tail adaptor rings, and a trimeric central tail fiber gp56 (needle) is anchored within the nozzle lumen, as described below. Portal protomer contacts with neck adaptor (Fig. S7). There are following types of contacts: a contact of portal stem domain with neck adaptor hanger; a contact between adaptor spine Ser190 and portal clip domain Lys409. The latter interface appears relatively flexible, suggesting that it may contribute to the reorientation of the tail body following DNA ejection, as discussed above.

The portal protein gp72 has a canonical fold comprising six domains: the stem and the clip domains, the crown and the wing domains, the barrel domain and the tunnel loop. The barrel consists of long α-helices that extend into the capsid interior. In the ejected state the density map allowed us to build a reliable model of the barrel domain spanning Pro610-Val696 residues, while only putative alpha helical overall chain trace in intact state. Density corresponding to the tunnel loop was partially resolved in both states. The differences between tunnel loops in intact and ejected states are laying within 1 Å of RMSD (Fig. 5B). The lumen of the portal complex is filled with a cylindrical density, which appears unstructured in the cryoEM density maps. This density represents the dsDNA and it is absent from the density map of ejected virions. Several regions of gp72 remain unresolved. The atomic model begins at Ala16, and residue range Arg501 - Asn527 lacks the density. The latter is enriched with arginines (its sequence is RRNHAVVINRDDRQRRQTIVLNAEGDN), suggesting this region to be involved in interactions with DNA.

Two dodecameric rings of adaptor proteins, gp64 and gp62, are attached to the portal and adopt almost identical folds (Fig. 5A), with pairwise Cα RMSD values ranging from <1 to 8 Å across separate domains (Fig. S3E). This fold is conserved among podoviruses and observed in phages as T7, Sf6, GP4, Pam1, ΦcrAss001 (Guo et al. 2014; F. Li et al. 2022; Zheng et al. 2023; J.-T. Zhang et al. 2022; Bayfield et al. 2023). The number of adaptor rings varies between phages, ranging from one to three. The low sequence identity between gp64 and gp62 (18%) suggests that gene duplication event giving rise to the second adaptor ring occurred early in evolution.

The gp62 adaptor performs a symmetry transition between the C12 symmetry of the gp64 ring and the C6 nozzle protein gp57. The differences between subunits are localized in regions Glu138-Val151 and Asp173-Ala178 which are involved in contact with the nozzle and lateral tail spikes. The RMSD between neighboring gp62 chains ranges from <1 to 3 Å. Within the lumen of the adaptor assembly, we observe diffuse, unstructured density that cannot be unambiguously assigned to either DNA or protein. This density is absent in the reconstruction of ejected virions, suggesting that the corresponding components are released together with the genomic DNA (Fig. 5B).

The nozzle protein gp57 assembles as a hexamer at the distal end of the tail following the gp62 adapter. The lumen of the gp57 is occupied with a trimer of a needle protein gp56. Comparison of intact and ejected virions reveals only minor changes in the positions of the nozzle tip domains that contact the needle, which is released upon DNA release. Apart from this small change and a minor transition in the portal barrel region, no substantial conformational changes are detected in the phi24B virion during genome release. This observation indicates that the portal and tail assembly behaves largely as a rigid body throughout DNA ejection.

The architecture of the phi24B portal and tail complex closely resembles that of *R. solanacearum* phage GP4 (identity and E-values: 27% and 1.7e-54 for portal, 23% and no E-value for neck adaptor, 33% and 4.7e-24 for tail adaptor, 29% and 2.9e-53 for nozzle). In contrast, the tail fibers of the two phages possess absolutely different fold and oligomerization. These observations suggest that the observed tail architecture represents a conserved structural core that allows phage adaptation through extensive modification or replacement of host-binding components.

#### The structure of the tail fibers

Cryo-EM reconstruction of the phi24B virion reveals two types of tail fibers (Fig. 6 A,B). Trimers of the gp61 protein form six lateral tail fibers, with their N-terminal domains attached to the tail and the C-terminal domains oriented toward the capsid. The axial fiber (tail needle) is formed with a trimer of the gp56, with its N-terminal domain anchored within the lumen of the nozzle formed by gp57 protein and C-terminal domain protruding from the tail via a long, flexible fiber.

**Figure 6:**
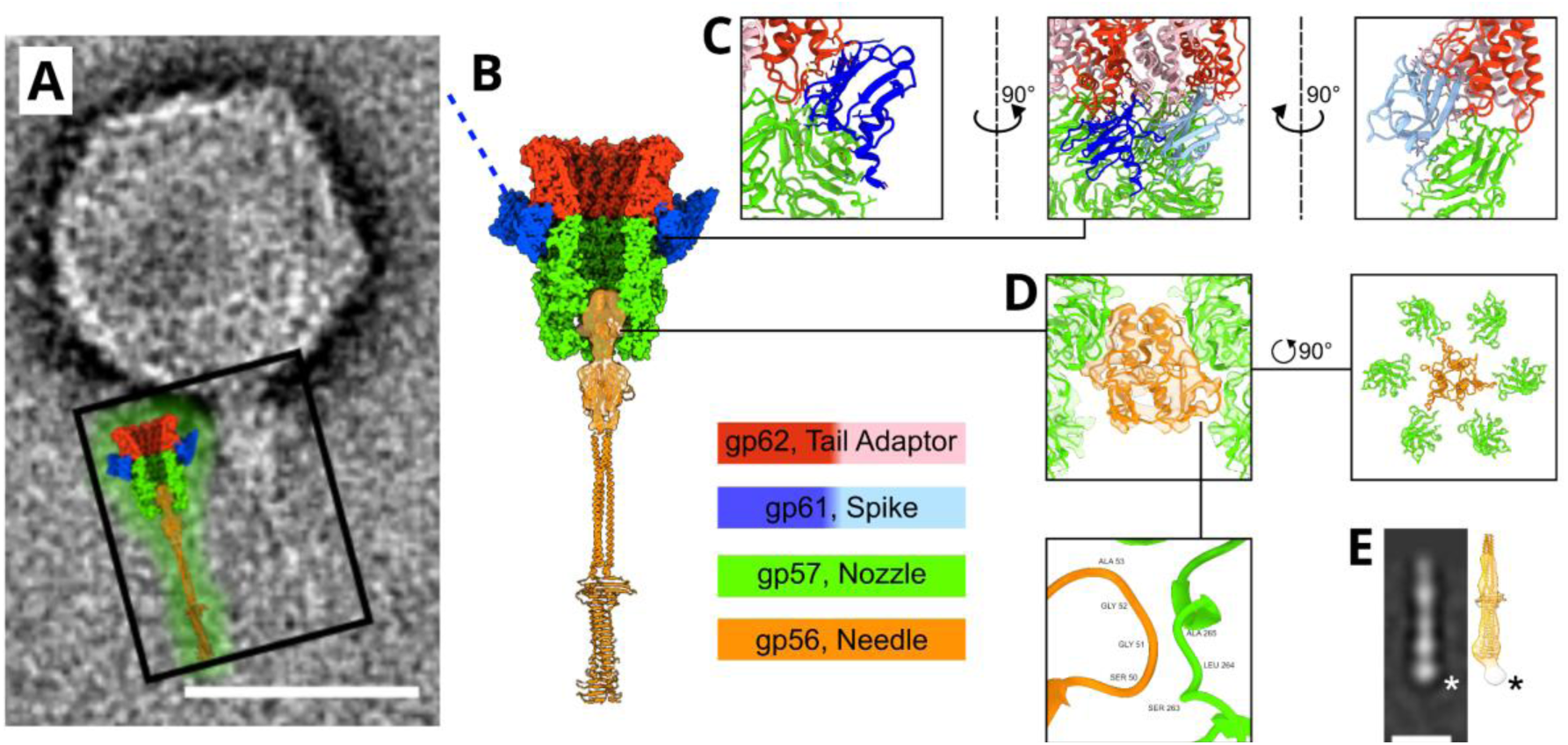
Structure of the phi24B tail fibers. A - CryoEM image illustrating the morphology of the tail spike gp61 and central needle gp56. The needle is visible as a long, flexible filament extending from the nozzle at the distal end of the tail. B - Molecular model of the gp56 needle predicted by AlphaFold3 and fitted into the cryo-EM density map. The needle is shown as a ribbon model, the corresponding density map is displayed semi-transparent, and the surrounding tail proteins are shown as surfaces. C - docking of the pair of trimer on tail body surface. D - Anchoring of the trimeric N-terminal domains of the needle between six nozzle tip domains at the distal end of the tail. The density map is shown semi-transparent. E - 2D class average from negative-stain electron microscopy showing the morphology of the distal end of the needle. The AlphaFold-predicted C-terminal β-prism domain of gp56 is fitted into the three-dimensional density map. An asterisk marks additional density at the needle tip, potentially corresponding to an additional protein.

We were able to model 141 of 645 residues of gp61 based on the cryo-EM density map. Within each trimeric spike, individual subunits are anchored to the adapter protein gp62 via a N-terminal β-sandwich domain. The β-sandwich domains are bound to the C-terminal alpha-domain of gp62 with charged interaction formed by gp61 Lys65, Lys67, as well as hydrogen bonds in the vicinity of Gln26. As there are a total of twelve gp62, only twelve N-terminal domains of gp61 are able to bind to them, thus in each trimer only two N-terminal domains are bound to the tail while the rest one is floating (Fig. 6B). This configuration leads to a fixed orientation of the fibers with the distal tip pointed toward the capsid. Beginning at residue 100, the spike transitions into a trimeric coiled coil typical for phage tail fibers, particularly phi24B close structural relative GP4 (Zheng et al. 2023).

Despite the expected flexibility of such coiled-coil fibers, a substantial portion of the fiber of ∼20 nm is visible in the cryo-EM density map. This suggests that its mobility may be restrained, possibly through interactions between the distal C-terminal region and the capsid surface. The C-terminal domain is visualized in two-dimensional class averages but cannot be resolved in the three-dimensional reconstruction. AlphaFold predictions likewise did not yield a confident structural model for this region, thus its interaction with the capsid remains unclear. We do not observe any differences in gp61 between the intact and ejected states of phi24B.

The needle protein gp56 exhibits substantial conformational flexibility. Individual cryo-EM micrographs of vitrified virions, as well as TEM images of negatively stained samples (Fig. 6A) show gp56 as a long, flexible axial fiber protruding from the virion, with an approximate length of 25 nm (Fig. 6A). Three-dimensional reconstruction was possible only for the N-terminal fragment of the needle, which is anchored within the gp57 nozzle (Fig. 6C). The remaining portion of the fiber is too flexible to be resolved and, owing to its relatively low molecular mass and thus limited signal-to-noise ratio, could not be reconstructed using conventional cryo-EM approaches such as local refinement.

The axial fiber protein N-terminal domain itself also exhibits mobility, resulting in reduced local resolution in this region together with the adjacent nozzle tip domains. At the achieved resolution of 10 Å, only the tertiary structure of the AlphaFold-predicted model could be fitted, with minimal adjustments applied to avoid overfitting (Fig. 6C). Nevertheless, the N-terminal domain could be unambiguously placed into the density map (Fig. 6C). The tail needle N-terminal domain contains extended flexible loops that protrude from the globular core and contact three of the six nozzle tip domains. These interactions occur in the vicinity of the gp57 loop spanning Arg47–Leu57 (Fig. 6D). Although the resolution does not permit visualization of side chains, the interacting loops contain charged residues that could support salt bridges and hydrogen-bond formation. The gp56 density is absent in the density maps of the ejected state of phi24B, which is consistent with the needle detachment upon DNA release.

Residues Val167–Met260 of the needle form a trimeric coiled-coil segment that grants the fiber its flexibility. The remaining C-terminal region assembles into a trimeric β-prism located at the distal tip of the needle. Owing to the low signal-to-noise ratio, we were unable to reconstruct this C-terminal domain from cryo-EM data; however, clean 2D class averages as well as a low-resolution 3D reconstruction was obtained from negatively stained samples (Fig. 6E). Rigid-body fitting of the C-terminal needle model into this map allowed us to visualize additional globular density at the distal tip. We propose that this density corresponds to a receptor-binding protein associated with the needle.

The molecular model of gp56 tail needle was not deposited in the Protein Data Bank, as it is largely based on AlphaFold prediction rather than cryoEM experiment; this model is provided as Supplementary Data.

## Discussion

Bacteriophage phi24B is one of the most dangerous viral agents associated with bacterial pathogenicity, which is responsible for conversion of some E. coli having moderate pathogenic potential into the causative agents of severe life-threatening infection. The type of the conversion by phi24B which may be rather termed lytic than lysogenic conversion, makes the in vivo toxin production not only dependent on lysogen induction frequency but, at least in theory, also on lytic multiplication of the virus on commensal E. coli populations in the patient’s gut. The lytic cycle of most of the coliphages is strongly inhibited by physico-chemical conditions of the human gut, structural features of phi24B described here may contribute to its ability to replicate and express toxin genes under such conditions, enabling efficient lytic replication and toxin gene expression even within the challenging environment of the human intestine.

From the perspective of genetic organization this restriction appears to be significantly relaxed in diarrhea (A. Letarov and Kulikov 2009). In addition, the lysogenization of new E. coli strains by this phage may lead to the emergence of novel EHEC strains with distinct epidemic potential. Therefore, structural properties of the phi24B virion, including the elements involved in the host cell recognition and interaction with the environment, are not only of biological interest, but may have direct practical implications.

Genomically, phage phi24B is a lambdoid virus sharing with the classical bacteriophage λ high similarity of genome organization and related lysogeny control region (Smith et al. 2012). Аt the same time, the structure of the phi24B virion is completely distinct from all the lambdoid phages studied previously. The closest structural homolog is Ralstonia phage GP4, whose virion core - comprising the capsid and tubular tail - is nearly identical to that of phi24B despite limited sequence similarity (for example, ∼33% amino acid identity between the MCPs of the two phages). However, GP4 is equipped with a very complex and unusual for tailed phages set of fibers (Zheng et al. 2023), while phi24B carries a more typical complex of six tail fibers and a needle.

The structural similarity between phage Gp4 and phi24B suggests that this virion architecture may represent an evolutionarily conserved module encompassing both capsid and tail genes. Notably, this structural module encodes a T=9 capsid which is larger compared to the more common T=7 capsids found in most lambdoid phages (Essus et al. 2023; Wang et al. 2022), therefore allowing phi24B to carry larger genome (57.7 kbp in phi24B compared to 48.7 kbp in λ).

Structural similarities are also observed in more distantly related viruses, such as the cyanophage Pam1 (J.-T. Zhang et al. 2022). The composition, oligomerization, and spatial arrangement of the phi24B cementing capsid protein gp67 closely resemble those of the Pam1 cementing protein gp6 (Cα RMSD ∼4.8 Å across all 124 aligned pairs). In PAM1, it has been proposed that the cementing protein may contribute to host recognition, as it shares structural similarity with the terminal domains of tail spike proteins. In light of this observation, we propose that the gp67 “tulip” assemblies in phi24B may likewise participate in host interactions. Such a function could be advantageous given the relatively large capsid (∼74 nm), which may shield the short tail when the virion approaches the host in a head-first orientation. In GP4, the cementing protein is similar. However, its N-terminal arm is positioned within the β-sandwich of the monomer, thereafter it lacks “hugging” contacts between neighboring monomers within the tulip.

Although the core virion structure of phi24B closely resembles that of Ralstonia phage GP4, the peripheral elements differ substantially. Whereas GP4 possesses three distinct types of tail fibers (Zheng et al. 2023), phi24B has only one type of the lateral fibers formed by the trimers of gp61 protein. This protein is highly conserved among phi24B-related viruses and was previously supposed to be responsible for the binding of the terminal phi24B receptor outer membrane BamA (Smith et al. 2007). Our cryoEM data resolve only the N-terminal part of gp61 suggesting the remainder of the protein is flexible. This finding is consistent with AlphaFold predictions, a very long collagen-like shaft (residues 243-567), interrupted by a short coiled-coil motif in the middle, which separates the N-terminal α-helical coiled coil rod and relatively small C-terminal globular domain (residues 568 - 644) potentially involved in the recognition of a receptor. Such an arrangement makes it unlikely that the conformational signal from the binding of the gp61 C-terminus to any host cell surface structure could be transmitted through the flexible collagen-like fiber to trigger the DNA release from the phage. Notably, sequences closely related to the collagen-like shaft and C-terminal domain of gp61 are also found in the lateral tail fibers of bacteriophage λ_2B8, initially obtained by prophage induction from children feces E. coli isolate 2B8 (Mathieu et al. 2020). In λ_2B8 these lateral fibers do not recognize the BamA protein of E. coli MG1655 (Kuznetsov et al. 2025) indirectly supporting the conclusion that in phi24B the closely related C-terminal domain of gp61 is not involved in BamA binding.

The central fiber (needle), formed by a trimer of gp56, therefore represents a plausible candidate for involvement in terminal receptor recognition. The needle is inserted into the nozzle (Fig. 6), and we have resolved the interactions between its N-terminal part and three of six surrounding nozzle tip domains. Although the map resolution of the N-terminal region was not sufficient for quantitative assessment of binding, the interface appears limited, involving only 2–3 residues per monomer, suggesting a relatively weak interaction. Therefore, we propose that the detachment of the needle may directly follow its binding to the receptor and trigger the release of the virion-encapsidated ejection protein gp47 and genomic DNA.

The distal part of the needle is missing from our reconstruction due to its high flexibility. Nevertheless, the needle length and overall shape are in perfect agreement with the AlphaFold predicted model featuring high confidence score especially at the C-terminal β-helical prism. The relatively simple surface of this domain makes it unlikely to mediate specific interactions with the extracellular loops of the BamA receptor.

However, low-resolution reconstruction of the distal needle tip from negative-stain transmission electron microscopy revealed an additional small, slightly asymmetric density (Fig. 6E). This feature is broadly consistent in size with the predicted heterodimer of gp54 and gp55, which are positioned downstream of the needle protein gp56. The larger protein, gp54 (204 amino acids), was detected in structural proteome at low estimated abundance, whereas gp55 (75 amino acids) was not detected, potentially due to its small size and low copy number. These observations raise the possibility that gp54, alone or in complex with gp55, associates with the distal end of the tail needle and may participate in BamA receptor recognition. This interpretation is consistent with previous bioinformatic predictions (Matusiak et al. 2025). However, further experimental studies are required to confirm the identity of the main phi24B RBP.

Bacteriophages may interact not only with bacterial surfaces, but also with components of their surrounding environment. For example, bacteriophage T4 was shown to bind mucin (Barr et al. 2015). Here, we demonstrate the conserved phi24B protein gp84, previously identified as carbohydrate esterase related to the E. coli mucin deacetylase NanS, is in fact a structural capsid decorating protein, bound to the central capsomers of the T=9 capsid facets (Fig. 4). Notably, gp84 (or at least its N-terminal capsid binding domain) interact with the capsomers in the form of hexamer, in contrast to most decorating proteins occupying local sixfold axes, which typically bind as monomers (for example, Hoc in T4-related phages (Rao et al. 2023), pb10 in T5-related viruses (Vernhes et al. 2017; Ayala et al. 2023)). An additional striking feature of gp84 is its proteolytical self-processing following phage release from the host cell (Fig. 1). In addition to the previously described cleavage between the catalytic domain and C-terminal lectin-fold domains (Barr et al. 2015), our data suggest the presence of a second autoprocessing site located between N-terminal capsid binding domain (previously referred to as unknown function domain DUF 1737) and the NanS-related catalytical domain.

The DUF1737 domain, which we demonstrate to bind capsomers on the virion exterior, is encoded not only in Enterobacteriaceae but also across diverse taxa within the phylum Pseudomonadota (including Moraxellales, Rhodobacteriales, and Rhizobiaceae) as well as in Actinomycetota. This broad distribution suggests that phages with particle architectures similar to phi24B are widespread and infect phylogenetically diverse hosts, pointing to an ancient evolutionary origin of DUF1737-containing capsid-decorating proteins. In addition, it can be hypothesized that NanS-p family esterase genes, to which gene 84 of phi24B belongs (Franke et al. 2020), may have originated in phages rather than in cellular genomes (Pascal et al. 2023). However, testing this hypothesis requires an in-depth analysis of the prevalence of the CDSs encoding SASA and DUF1737 domains in bacterial genomes and their relation to prophage sequences,which is beyond the scope of this study.

The biological significance of the delayed proteolytic maturation of gp84 remains unclear. It is possible that gp84 at the capsid surface helps the phage to concentrate at the mucus and, maybe, to overcome the infection inhibition that mucin exerts to some phages (Green et al. 2021). Then the release of mucin-binding phi24B may facilitate the phage spread out of the primary locus of its propagation. Elucidating the functional role of gp84 will require further experimental investigation, including analysis of mutants with disrupted processing sites.

In summary, we may say that being at first glance a trivial podovirus, the bacteriophage phi24B turned out to have a plethora of unusual structural properties highlighting several directions of possible experimental and theoretical research. Among the questions streaming from our structural data are:

1. What is the evolutionary origin of the GP4–phi24B structural module, and why is it rare among characterized phages despite being conserved across distantly related viruses?
2. What is the biological function of the conserved lateral fibers (gp61), and how do they contribute to phi24B-related phages spread in natural conditions and the emergence of new EHEC strains?
3. How does the complex proteolytic maturation of the decorating protein gp84 contribute to phage fitness?
4. If receptor engagement is mediated primarily by the tail needle, how is stable attachment to the host cell maintained following needle dissociation?
5. How is the exceptionally large ejection protein gp47 (>3,000 amino acids) organized within the capsid?

Addressing these questions will provide deeper understanding of the biology and ecology of the most widely spread Stx-converting phages and their role in pathogen evolution and epidemiology.

## Materials and Methods

### Phage purification

#### Phage and bacterial strains and their propagation

The bacteriophages phi24B:Cat (Allison et al. 2003) was a king gift by professor Heather Allison from Liverpool University, UK. For the phage plaque assay bacterial host strain E. coli K-12 MG1655 was used. The lysogen E. coli 4sR (phi24B:Cat) was obtained by us earlier (Golomidova et al. 2021) by lysogenizing the rough derivative of environmental E. coli isolate 4s. For both host and phage propagation the LB medium (Trypton (Amresco) 10g, Yeast extract (Amresco) 5g, NaCl 10g, deionized water up to 1L). LB broth was supplemented by 15 g/L of the Bacto-Agar for the plates and with 6 g/L1 of the Bacto-Agar for top agar for bacteriophage titration.

The phage titration was performed using the conventional double-layer method but in order to improve the plaques visibility the top-agar was supplemented by a sub-inhibitory concentration (2,5μg/mL) of chloramphenicol. For host lawn 1 mL of overnight culture and 2 mL of soft LB agar was used with addition of CaCl2 and MgSO4, final concentration 10mM each.

#### Phage stock preparation and purification

According to the developed method (Kuznetsov et al. 2024) pure and concentrated virion suspension was obtained. Prophage induction in lysogenic cells derived from the 4sR strain (Golomidova et al. 2021) was performed as follows. Mitomycin C was added to a final concentration of 1 μg/mL, and MgSO₄ to a final concentration of 10 mM, to a liquid culture of 4sR (phi24B:Cat) lysogens grown in LB to an OD₆₀₀ = 0.1. The culture was then incubated overnight at +37°C with shaking at 250 rpm under low aeration conditions (using 300 mL Erlenmeyer flasks filled with 250 mL of culture).

Chloroform was added to the obtained phage lysates, and incubation was continued under the conditions specified above for 30 minutes, after which the lysates were clarified by centrifugation at 15,000×g for 20 minutes. All subsequent ultracentrifugation steps were carried out at 75,000×g, with maximum acceleration and deceleration rates, at a temperature of +20°C for one hour. The clarified lysates were first concentrated by ultracentrifugation using a Beckman 45Ti fixed-angle rotor. The concentrated samples were then resuspended in SM buffer (8 mM MgSO₄, 100 mM NaCl, 50 mM Tris-HCl pH 7.5) and layered onto a stepwise sucrose density-viscosity gradient (20%-30%-40%-50%-60%) prepared in buffer (50 mM Tris-HCl, pH 7.2, 50 mM NaCl). This was followed by ultracentrifugation in a Beckman SW 50.1 swing-bucket rotor. After centrifugation, purified phage particles were collected from the interface between the 40% and 50% sucrose layers. The preparation was then dialyzed against the SM buffer and subjected to an additional concentration step using Freon. The purified preparation was diluted in SM buffer and layered onto a cushion of Freon CFC-113 (density 1.56 g/cm³), followed by centrifugation under the same parameters in the Beckman SW 50.1 rotor. The biological titer of the phage in the concentrate prepared this way reached up to 6.5 × 10¹² PFU/mL.

### Phage structural proteome analysis

#### Trypsinolysis in gel

The protein bands after 1D-PAGE were excised and washed with 1 ml of 100 mM NH4HCO3 and 50 % ACN mixture (v/v) at 50oC with constant shaking until the piece of gel becomes transparent. Protein cysteine bonds were reduced with 10mM DTT in 50 mM NH4HCO3 for 30 min at 56 °C and alkylated with 55 mM iodoacetamide in the dark at RT for 30 min. The step with adding DTT was repeated. Then gel pieces were dehydrated with 100 μl of acetonitrile, air-dried and treated by 20 μl of 15 mg/mL solution of trypsin (Trypsin Gold, Mass Spectrometry Grade, Promega) in 50 mM ammonium bicarbonate for 16 h at 37oC. Peptides were extracted with 50 μl of 50% acetonitrile (ACN)/ 5% formic acid (FA) solution for 30 min with sonication. This extraction procedure was repeated 3 times. Peptides were dried under vacuum and redissolved in 5% ACN with 0.1% FA solution prior to LC-MS/MS analysis.

#### Trypsinolysis in solution

Samples were incubated with ProteinSafe™ Protease Inhibitor Cocktail (TransGen, China) and benzonase (diaGene, Russia; 5 U per 100 µL of sample) for 10 min at 37 °C, followed by incubation with 4% SDS for 30 min at 60 °C. Phage particles were then lysed by sonication at 40% amplitude using a pulse regime of 15 s on and 30 s off for a total processing time of 3 min, with the temperature maintained at 4 °C. Lysates were clarified by centrifugation at 15,000 × g for 10 min, and the supernatants were transferred to fresh tubes.

Proteins were precipitated using a chloroform–methanol extraction protocol. Briefly, 400 µL of methanol was added to 100 µL of protein sample in a 1.5 mL microcentrifuge tube and mixed thoroughly by vortexing. Subsequently, 100 µL of chloroform was added, followed by additional vortexing. Next, 300 µL of Milli-Q water was added, and the mixture was vortexed and centrifuged at 14,000 × g for 1 min. The upper aqueous phase was carefully removed. Proteins were then precipitated by addition of 400 µL of methanol, followed by vortexing and centrifugation at 20,000 × g for 5 min. The supernatant was discarded, and the resulting protein pellet was dried under vacuum.

Precipitated proteins were denatured for 30 min in a buffer containing 8 M urea, 2 M thiourea, and 10 mM Tris (pH 8.0). Protein concentration was determined using a Bradford assay, and 200 µg of protein from each sample was used for subsequent analysis. Disulfide bonds were reduced with 15 mM dithiothreitol (DTT) at 50 °C for 30 min and alkylated with 30 mM iodoacetamide for 30 min at room temperature in the dark. The reduction step with DTT was then repeated, after which the reaction mixture was diluted sixfold with 50 mM Tris–HCl (pH 8.0).

Proteolytic digestion was performed using Trypsin Gold (mass spectrometry grade; Promega) at an enzyme-to-protein ratio of 1:50 (w/w) overnight at 37 °C. Digestion was terminated by acidification with trifluoroacetic acid (TFA) to a final concentration of approximately 1%. The resulting peptide mixture was desalted using Copure C18A solid-phase extraction (SPE) cartridges (100 mg mL⁻¹; Biocomma), dried under vacuum, and reconstituted in 5% acetonitrile (ACN) containing 0.1% formic acid (FA) prior to LC–MS/MS analysis.

#### LC–MS/MS analysis

LC–MS/MS analysis of peptide samples was performed using an Orbitrap Exploris 480 mass spectrometer (Thermo Scientific, USA) coupled to an Ultimate 3000 RSLCnano chromatographic system (Dionex, USA) via a Nanospray Flex ion source (Thermo Scientific, USA).

Peptide samples were resuspended in 35 µL of 5% acetonitrile (HPLC grade) in water (LC–MS grade) containing 0.1% (v/v) formic acid. Chromatographic separation was performed by reversed-phase liquid chromatography using a C18 PepMap 100 trap column (C18 sorbent, 5 mm length, 300 µm inner diameter, 5 µm particle size, 100 Å pore size; Thermo Scientific, USA) coupled to a Peaky-C18 analytical capillary column (C18 sorbent, 50 cm length, 75 µm inner diameter, 1.9 µm particle size; Molekta, Russia).

For each analysis, 5 µL of sample was loaded onto the trap column using 5% acetonitrile (HPLC grade) in water (LC–MS grade) with 0.1% (v/v) formic acid at a flow rate of 10 µL min⁻¹ for 12 min. The trap column was then switched in line with the analytical column. Peptides were eluted using a binary solvent system consisting of solvent A (5% acetonitrile in water, 0.1% formic acid, v/v) and solvent B (79.9% acetonitrile, 20% water, 0.1% formic acid, v/v).

The gradient was applied at a flow rate of 250 nL min⁻¹ as follows: solvent B was increased from 5% to 10% over 3 min, from 10% to 55% over 54 min, and from 55% to 65% over 3 min. After peptide elution, the system was washed with 100% solvent B for 2 min and subsequently re-equilibrated with 5% solvent B for 15 min.

The electrospray ionization source was operated at a spray voltage of 2,700 V, and the capillary temperature was set to 275 °C. The mass spectrometer was operated in data-dependent acquisition (DDA) mode. In each acquisition cycle of 1 s, a full MS survey scan was recorded, followed by automatic selection of precursor ions for fragmentation and acquisition of MS/MS spectra. Precursor ion selection was performed automatically, and dynamic exclusion was enabled to prevent repeated fragmentation of ions for which MS/MS spectra had already been acquired.

Full MS spectra were acquired at a resolution of 60,000 over an m/z range of 200–1,500, with the automatic gain control (AGC) setting set to “Standard” and the maximum injection time set to “Auto”. The Funnel RF level was set to 50. MS/MS spectra were acquired at a resolution of 15,000 with the AGC setting “Standard” and an automatic injection time. Fragmentation was performed using a normalized collision energy of 30, with an isolation window of 1.4 m/z.

Quantitative analysis of MS/MS data was performed using MaxQuant software (Cox and Mann 2008), with peptide identification carried out by the Andromeda search engine against the Bacteriophage phi24B genome annotation database.

The abundance of the proteins in the proteome was estimated based on the detected intensity (peptide counts) in the batch hydrolysis experiment with the reference to the proteins the copy number of which can be derived from the structural data. We calculated estimated abundances using the lateral fiber (gp61), the nozzle protein (gp57), the needle protein (gp56) and the portal protein (gp72) as the references (Table S1, sheet 2).

The following formula was used for the estimation:

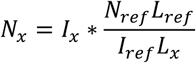

where *N*_*x*_ – copy number of the protein x per virion; *L*_*x*_ – the length of the polypeptide chain of the protein x, a.a.; *I*_*x*_ – signal intensity of the protein x in the batch hydrolysis experiment; *N*_*ref*_ – structure-derived copy number of the reference protein; *L*_*ref*_ and *I*_*ref*_ – length and intensity of the reference protein, respectively.

#### Native peptides sample preparation

To samples were added ProteinSafe™ Protease Inhibitor Cocktail (TransGen, China) and benzonase (diaGene, Russia; 5 U per 100 µL of sample) to inhibit proteolytic degradation and digest nucleic acids, respectively. Phage particles were then lysed by incubating at 95°C for 5 minutes and subsequent sonication at 40% amplitude using a pulse regime of 15 s on and 30 s off for a total processing time of 3 min, with the temperature maintained at 4 °C. Lysates were clarified by centrifugation at 15,000 × g for 10 min, and the supernatants were transferred to fresh tubes.

Following sonication, the lysate was treated with a high-concentration chaotropic agent to complete protein denaturation and solubilization. A volume of 8 M guanidine hydrochloride (GuHCl) solution was added to the sample to achieve a final concentration of 6 M GuHCl. A protease inhibitor cocktail was added again to maintain a 1x final concentration throughout the procedure. The mixture was then incubated for 30 minutes at room temperature with constant gentle mixing on a laboratory rotator.

To remove insoluble cellular debris, the sample was centrifuged at 20,000 g for 30 minutes at 4°C. The resulting supernatant, containing the solubilized proteins, was carefully collected. The resulting peptide mixture was desalted using Copure C18A solid-phase extraction (SPE) cartridges (100 mg mL⁻¹; Biocomma), dried under vacuum, and reconstituted in 5% acetonitrile (ACN) containing 0.1% formic acid (FA) prior to LC–MS/MS analysis.

### Cryo-electron microscopy

#### Data collection

Cryo-electron microscopy was performed with Titan Krios 300 kV microscope equipped with a Gatan K3 camera. Samples were applied to the glow discharged Quantifoil R1.2/1.3 holey carbon support film and vitrified using a Vitrobot Mark IV (Thermo Fisher Scientific) at 100% humidity and 4 °C. In total, three distinct datasets were collected, two with pixel size 0.83 Å and one with pixel size 1.06 Å. Data was imaged with dose of 53e-/ Å2 and −2.5 micrometer defocus. A total of 8460 movies were collected. Three distinct datasets were combined together with Fourier crop to pixel size 1.07 Å in CryoSPARC software v 4.7.1 (Punjani et al. 2017). The overall data of 8,460 movies were used for subsequent processing.

#### Single particle data processing

Data processing was carried out with CryoSPARC software v 4.7.1. The overall data collection workflow is presented in Fig. S8. A total of 8,460 dose-weighted micrographs were subjected to patch-based motion correction and patch-based CTF estimation. The dataset was curated by removing low-quality micrographs, resulting in 7,957 micrographs used for particle picking.

Manual picking with a diameter of 745 Å was used to generate templates for automatic picking of capsid projections. Automatic picking was then performed based on two templates corresponding to intact and ejected states made with manual picking. Therefore, 52,535 particles for intact virions and 35,536 particles for ejected virions were collected. After 2D classification, subsets of 18,404 intact and 18,123 ejected particles were selected for whole-virion asymmetric (C1) reconstruction.

An initial model for refinement was generated as a Gaussian-smoothed icosahedron with a diameter of ∼75 nm. 3D Homogenous Refinement with forced icosahedral symmetry and initial lowpass resolution of 140 Å for isometric icosahedral capsids was an initial step for further refinements. The reconstructions reached resolutions of 3.8 Å for intact capsids and 4.0 Å for ejected capsids.

Symmetry expansion with icosahedral symmetry was applied to the particle sets to isolate specific capsid regions: edges, faces, faces focused on Decorating Capsid protein (DCP) and vertices. Particle centers were reoriented toward these regions using the volume alignment tools. As a result, we collected clean particle sets of regions of C5 vertices without tail (2,915,483 symmetry-expanded particles for intact and 1,958,961 for ejected states), C3 faces (3,148,455 symmetry-expanded particles for intact and 2,129,977 particles for ejected states) and C2 edges (3,149,952 symmetry-expanded particles for intact and 2,130,009 for ejected states). To identify tail-containing particles, 3D classification was performed on the vertex subset using a cylindrical mask to isolate the unique symmetry-breaking tail vertex. Particles assigned to this class were further processed by reorienting particle centers toward the tail region, followed by focused reconstructions of the tail body, neck, and portal barrel.

All focused reconstructions were refined using local refinement combined with both local and global CTF refinement. The overall resolution achieved was ∼2.6 Å for capsid regions and ∼3.0 Å for tail regions (Fig. S5). As a final step, map sharpening was performed using phenix.auto_sharpen for each focused reconstruction.

Particles for asymmetric whole-virion reconstructions were selected from the icosahedral capsid reconstruction set. Several iterative rounds of 2D classification with progressively increased maximum alignment resolution were performed to remove particles overlapping with carbon edges or affected by severe ice contamination. An initial C1 reconstruction was generated by homogeneous refinement using a low-pass filtered whole-virion map of phage GP4 (EMDB:36464) as a reference. A series of subsequent 3D classifications with a cylindrical mask encompassing the tail region, each followed by asymmetric refinement, enabled the selection of particle subsets with orientations yielding reconstructions in which the tail density was consistently localized at a single capsid vertex. At this stage, the density of the fivefold capsid vertex was well resolved, whereas the sixfold tail density appeared smeared due to symmetry mismatch. To address this, focused 3D classification was performed without altering particle coordinates, using a cylindrical mask surrounding the tail region. This yielded five classes in which both the fivefold capsid vertex and the sixfold tail density were resolved, corresponding to discrete rotational positions of the tail relative to the capsid arising from symmetry mismatch. The classes were related by rotations of approximately n × 72° (n = 0–4) around the tail axis. The corresponding 3D class volumes and particle orientations were then brought into a common reference frame using volume alignment tools. The updated particle sets were subsequently merged. This procedure yielded combined datasets of 18,404 particles for intact and 18,123 particles for ejected virions.

For intact and ejected virions, the corresponding particle sets were independently subjected to variability analysis in cluster mode. This analysis revealed two distinct tail conformations. The particle distribution between the two conformers was 9,656 and 8,748 particles for intact virions, and 10,407 and 7,716 particles for ejected virions. The resolutions of asymmetric whole-virion reconstructions were 5.6 Å and 5.7 Å for the intact state conformers and 5.8 Å, 6.3 Å for the ejected state conformers. For each conformer in both intact and ejected states, a soft-edged mask was generated and used for local asymmetric refinement. Each mask encompassed 12 portal protomers (excluding the barrel domains) and 10 surrounding MCP subunits derived from symmetrized maps. These masks were used to obtain focused reconstructions of the portal–capsid interface. The resulting resolutions were 4.2 Å and 4.4 Å for the intact state, and 4.1 Å and 4.4 Å for the ejected state.

For focused analysis of the tail tip binding site, particles from both conformations in the intact dataset were combined and recentered using volume alignment tools. Following 2D classification to remove low-quality particles (including overlapping particles, those near carbon support edges), 13,303 projections were selected for local C3 refinement using a soft-edged mask centered on the N-terminal region of the gp56 needle.

#### Transmission electron microscopy with negative staining

To visualize the C-terminal region of the tail needle, we performed transmission electron microscopy with negative staining. A 3 μl aliquot of the sample was applied to a glow-discharged amorphous carbon support film and stained with 2% uranyl acetate for 30 s. Images were acquired using a JEM-2100 transmission electron microscope (JEOL, Japan) operating at 200 kV, equipped with a DE-20 camera (Direct Electron, USA). A total of 480 micrographs were collected at a defocus of −1.8 μm and a pixel size of 1.4 Å.

Due to adhesion to the carbon substrate, most virions were oriented with the gp56 needle lying along the surface. Negative staining enhanced the signal-to-noise ratio of the small C-terminal region of gp56, which is not resolved in cryo-EM reconstructions. Manual picking of needle tip fragments yielded 2,245 particle projections. These were subjected to 2D classification, resulting in four classes with similar morphology. The selected particles were then used to generate a low-resolution three-dimensional reconstruction using a helical refinement algorithm in cryoSPARC, without imposing helical symmetry.

#### Model Building, Refinement, and Validation

Protein sequences corresponding to potential virion components (gp57, gp61, gp62, gp64, gp67, gp69, gp72 and gp84) were selected from the previously published phi24B genome (Smith et al. 2012). Initial structural models were generated using the AlphaFold2 implementation in ColabFold (Jumper et al. 2021; Mirdita et al. 2022). To determine individual density regions in reconstructed 3D volumes the segmentation of reconstructed 3D density maps was performed using the Segger watershed algorithm (Pintilie et al. 2010) implemented in UCSF ChimeraX (Goddard et al. 2018) as “Segment map” tool. The predicted models were fitted as rigid bodies into the segmented densities and showed model–map cross-correlation coefficients in the range of 0.5–0.7.

Model refinement was carried out iteratively. First, manual refinement was performed in ChimeraX using the ISOLDE molecular dynamics flexible fitting (MDFF) extension (Croll 2018), with predicted aligned error (PAE) matrices applied as restraints. Ambiguous or poorly resolved regions were adjusted or truncated in Coot (Emsley et al. 2010). Second, restraint parameters for real-space refinement were generated in ISOLDE using the “isolde write phenixRsrInput” command. The models were then refined using Phenix real_space_refinement (Afonine et al. 2018), applying restraints that limited large-scale movements while improving local geometry (bond lengths, angles, and stereochemistry).

Model validation was performed using MolProbity (Williams et al. 2018) within the Phenix suite (phenix.validation_cryoem) (Table S1). Models exhibiting suboptimal validation metrics or poor model–map agreement were iteratively corrected in ISOLDE and subjected to further refinement and validation. Final assemblies were generated using the sym command in ChimeraX. Cryo-EM density maps and atomic models were visualized using ChimeraX. A schematic representation of the phi24B genome region of interest was generated using DNA Features Viewer (https://github.com/Edinburgh-Genome-Foundry/DnaFeaturesViewer).

## Supporting information

Supplementary Data - Proteome

## Acknowledgments

We’re grateful to Prof. Heather Allison from the Liverpool University, UK, for sharing phage phi24B:Cat and useful discussion and also to Dr. Mikhail Shneider from Shemyakin and Ovchinnikov Institude of Bioorganic Chemisctry, Moscow, Russia for critical evaluation of our structural models and insightful suggestions which helped to significantly improve our results. We’re also grateful to Prof. Grzegorz Wegrzyn, Dr. Silwia Bloch and Dr. Bozena Nejman-Falenczyk from the University of Gdansk, Poland for their kind help in establishment of phi24B bacteriophage cultivation in our laboratory.

The research was funded by RSF 25-24-00110. CryoEM studies were carried out at the Shared Research Facility “Electron microscopy in life sciences” at Moscow State University (Unique Equipment “Three-dimensional electron microscopy and spectroscopy”) and at Kobilka Institute of Innovative Drug Discovery (KIIDD), School of Medicine, CUHK, China.

## Data availability

The asymmetric atomic models of φ24B capsids and tails in intact and ejected states have been deposited in the Protein Data Bank (PDB) under accession codes 9XLR, 9XLQ, 9XIW and 9XIV. Atomic models of capsid faces with and without the decorating capsid protein gp84 N-terminal domain, in both intact and ejected states, have been deposited under accession codes 9XLP, 9XLL, 9XLM and 9XLN.

The corresponding cryo-EM density maps have been deposited in the Electron Microscopy Data Bank (EMDB). For intact capsids and their components (vertices, faces with and without DCP, and edges), accession codes are EMD-66999, EMD-67000, EMD-67001, EMD-67002, EMD-67004, EMD-66996 and EMD-67003. Tail reconstructions for the intact state are available under EMD-66914, EMD-66919 and EMD-66920.

For ejected capsids and their components, maps are available under EMD-66992, EMD-66995, EMD-66993, EMD-66994, EMD-66997, EMD-66998 and EMD-66991, and for tail reconstructions under EMD-66916, EMD-66917 and EMD-66915. Asymmetric reconstructions of capsid units are available under EMD-67006 and EMD-67005, and of tails under EMD-66921 and EMD-66918, for intact and ejected states, respectively.

Cryo-EM reconstructions of whole virions in intact and ejected states, including both tail conformations, are deposited under accession codes EMD-69679, EMD-69695, EMD-69713 and EMD-69716. Focused reconstructions of capsid–tail interfaces are available under EMD-69717, EMD-69718, EMD-69719 and EMD-69720. The focused reconstruction of the N-terminal region of the gp56 tail needle has been deposited under accession code EMD-69734.

## Supplementary Materials

**Table S1.** Merged Maps and Models refinement statistics.

|  | Capsid<br>Asym-<br>etric<br>Unit,<br>Intact<br>State | Capsid<br>Asym-<br>etric<br>Unit,<br>Ejected<br>State | Tail,<br>Intact<br>State | Tail,<br>Ejecte<br>d State | Face<br>with<br>DCP,<br>Intact<br>State | Face<br>without<br>DCP,<br>Intact<br>State | Face<br>with<br>DCP,<br>Ejected<br>State | Face<br>without<br>DCP,<br>Ejected<br>State |
| --- | --- | --- | --- | --- | --- | --- | --- | --- |
| Density<br>map EMDB<br>code | EMD-67006 | EMD-67005 | EMD-66921 | EMD-66918 | EMD-67004 | EMD-66996 | EMD-66997 | EMD-66998 |
| Atomic<br>model PDB<br>code | 9XLR | 9XLQ | 9XIW | 9XIV | 9XLP | 9XLL | 9XLM | 9XLN |
| Map<br>resolution<br>(FSC 0.143<br>cut-off) | 2.4 | 2.4 | 2.9 | 3 | 2.7 | 2.9 | 2.9 | 2.9 |
| Map to<br>model<br>correlation<br>coefficient | 0.78 | 0.79 | 0.78 | 0.78 | 0.83 | 0.85 | 0.85 | 0.85 |
| Bond<br>lengths, Å | 0.004<br>(0) | 0.004<br>(0) | 0.007<br>(0) | 0.009<br>(0) | 0.004<br>(0) | 0.004<br>(0) | 0.004<br>(0) | 0.004<br>(0) |
| (# outliers<br>>4σ) | | | | | | | | |
| Bond angles, ° (# outliers<br>>4σ) | 0.926<br>(0) | 0.934<br>(0) | 0.95<br>(0) | 0.982<br>(0) | 1.081<br>(0) | 1.095<br>(0) | 1.087<br>(0) | 1.101<br>(0) |
| MolProbity Score | 1.22 | 1.18 | 1.05 | 1.25 | 1.15 | 1.13 | 1.14 | 1.12 |
| Clash score | 4.41 | 3.94 | 2.6 | 4.77 | 3.57 | 3.38 | 3.5 | 3.33 |
| CaBLAM outliers (%) | 0.64 | 0.76 | 0.63 | 0.64 | 0.64 | 0.34 | 0.68 | 0.38 |
| Rotamer outliers (%) | 0 | 0 | 0 | 0.08 | 0 | 0 | 0 | 0 |
| Ramachandran statistics (%) |  |  |  |  |  |  |  |  |
| Outliers | 0 | 0 | 0 | 0.03 | 0 | 0 | 0 | 0 |
| Allowed | 0.59 | 0.23 | 1.04 | 1.4 | 0.59 | 0.68 | 0.37 | 0.42 |
| Favored | 99.41 | 99.77 | 98.96 | 98.57 | 99.41 | 99.32 | 99.63 | 99.58 |
| Rama-Z score |  |  |  |  |  |  |  |  |
| whole | -1.42<br>(0.10) | -1.41<br>(0.10) | -1.52<br>(0.14) | -1.62<br>(0.14) | -1.37<br>(0.13) | -1.20<br>(0.14) | -1.44<br>(0.13) | -1.27<br>(0.14) |
| helix | -1.73<br>(0.15) | -1.73<br>(0.15) | -1.72<br>(0.13) | -1.55<br>(0.12) | -1.33<br>(0.19) | -1.24<br>(0.20) | -1.42<br>(0.19) | -1.26<br>(0.21) |
| sheet | -0.17<br>(0.13) | -0.14<br>(0.13) | -0.31<br>(0.20) | -0.59<br>(0.21) | -0.45<br>(0.17) | -0.25<br>(0.18) | -0.51<br>(0.17) | -0.34<br>(0.18) |
| loop | -1.02<br>(0.10) | -1.05<br>(0.10) | -0.49<br>(0.16) | -0.59<br>(0.16) | -0.87<br>(0.13) | -0.80<br>(0.14) | -0.87<br>(0.13) | -0.83<br>(0.13) |

**Figure S1.**
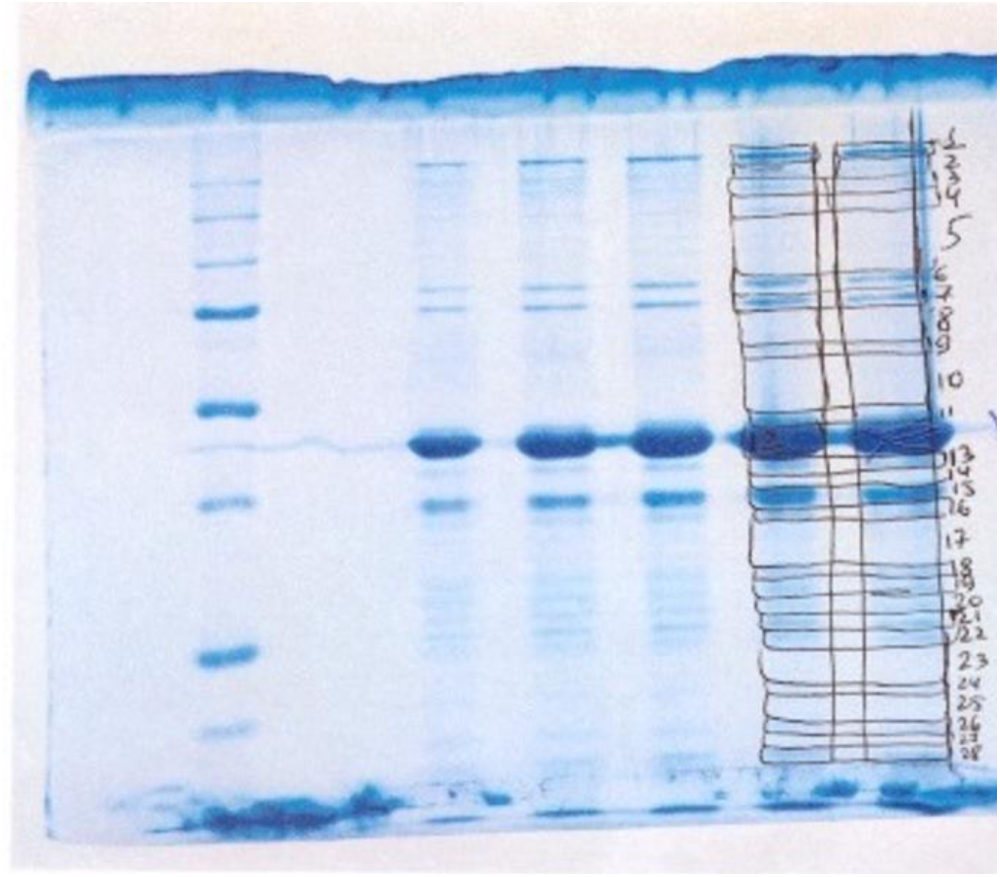
The bands of phi24B protein profile excised for proteins identification.

**Figure S2.**
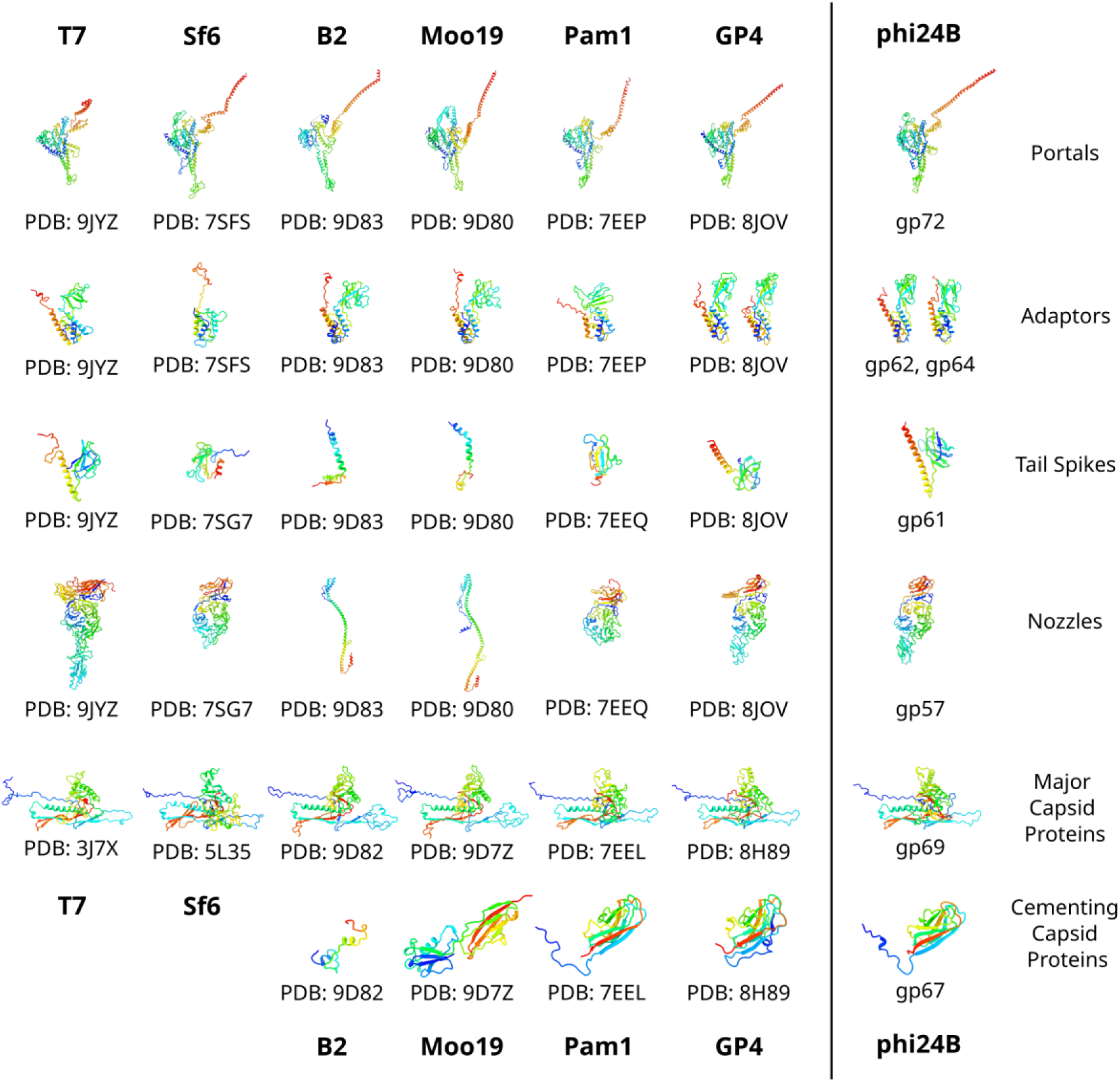
Comparison of structural components of phi24B with structurally relative phages. Molecules are colored by rainbow gradients from blue N-termini to red C-termini.

**Figure S3.**
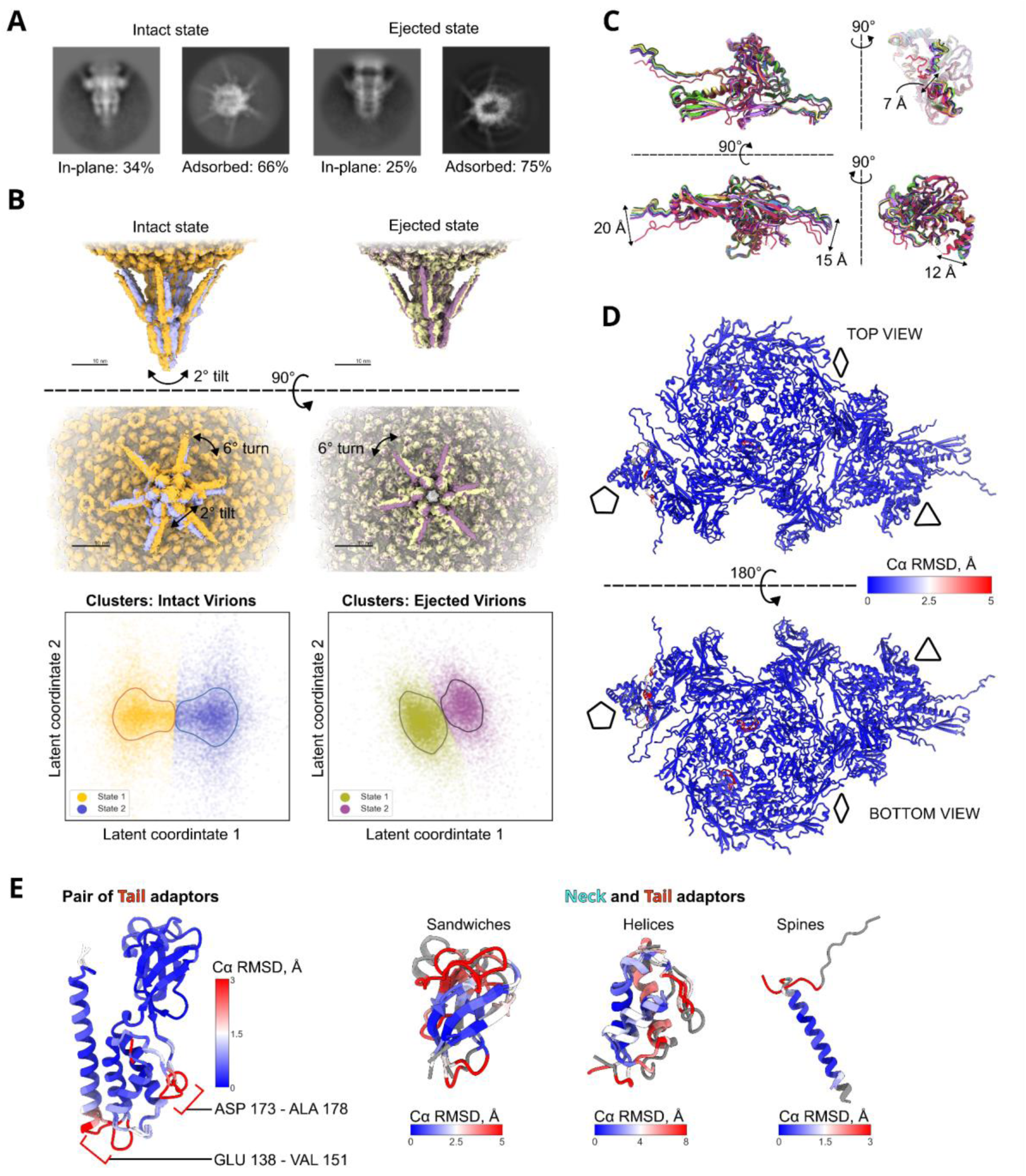
Variability analysis. A - Representative 2D classes and their relative proportions in tail directions distribution. B - Superposition of the two tail conformational states in intact and ejected virions, derived from clustering of latent components. C - Superposition of all nine MCP subunits within the asymmetric unit. D - RMSD between intact and ejected capsid asymmetric units. E - superimposition of pairs of adaptors colored by RMSD.

**Figure S4.**
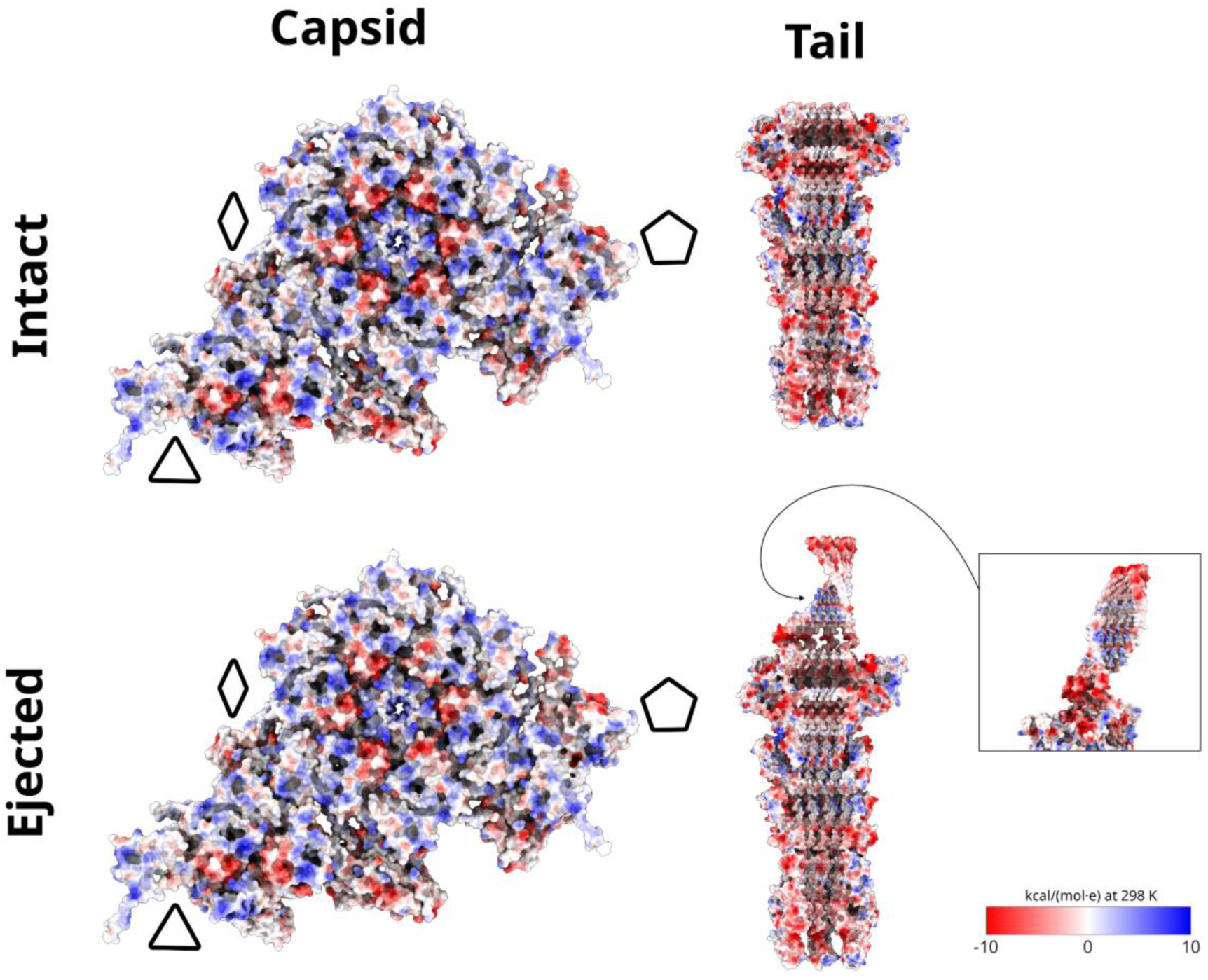
Analysis of Coulombic electrostatic potential. A - Electrostatic potential mapped onto capsid asymmetric units, viewed from the capsid interior. The pentagon, triangle, and rhombus denote fivefold, threefold, and twofold symmetry axes, respectively. B - Electrostatic potential of the tail assembly.

**Figure S5.**
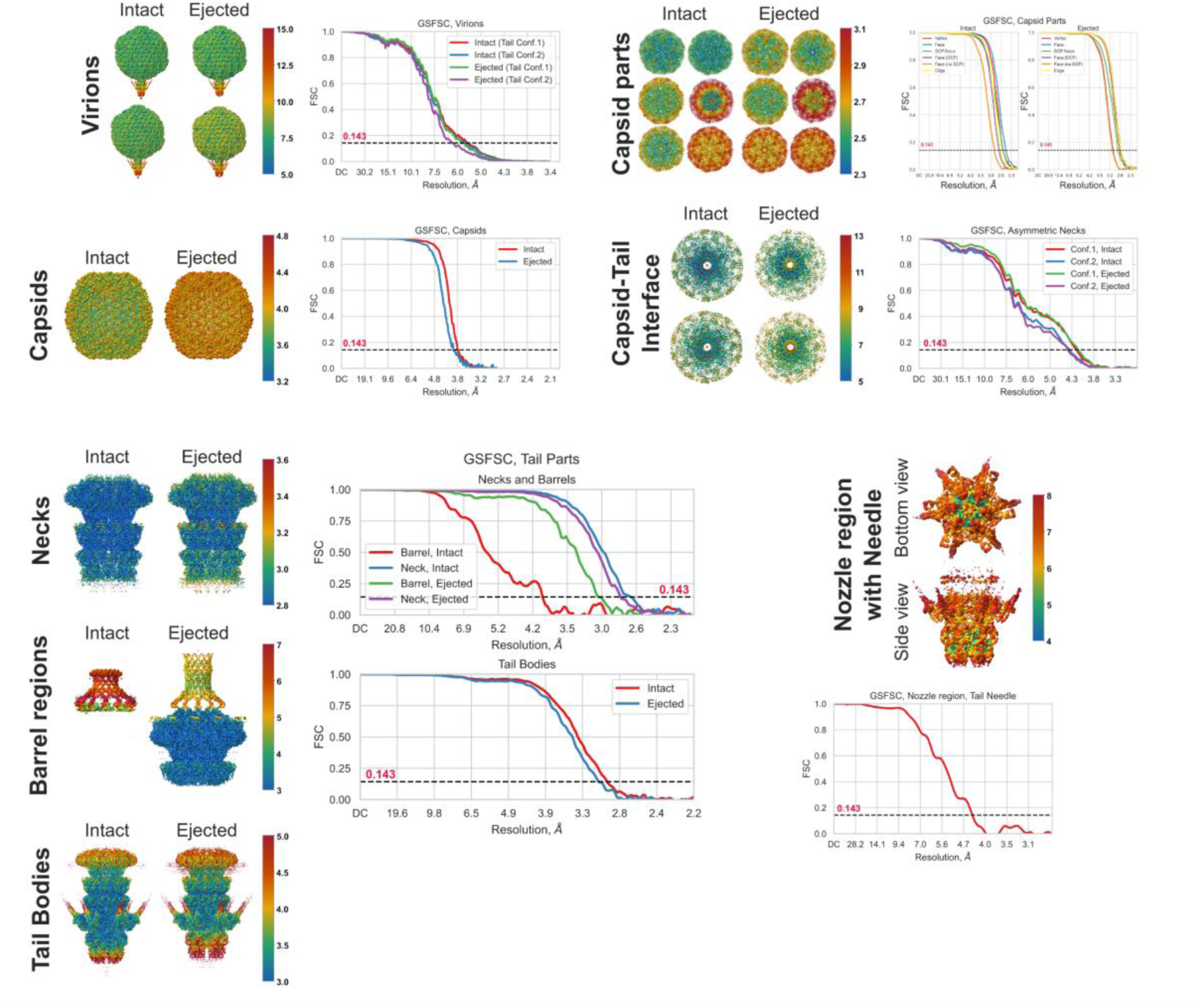
Local resolutions and GSFSC plots of obtained reconstructions of different parts of phi24B Intact and Ejected virions. (Tight curves from cryoSPARC)

**Figure S6.**
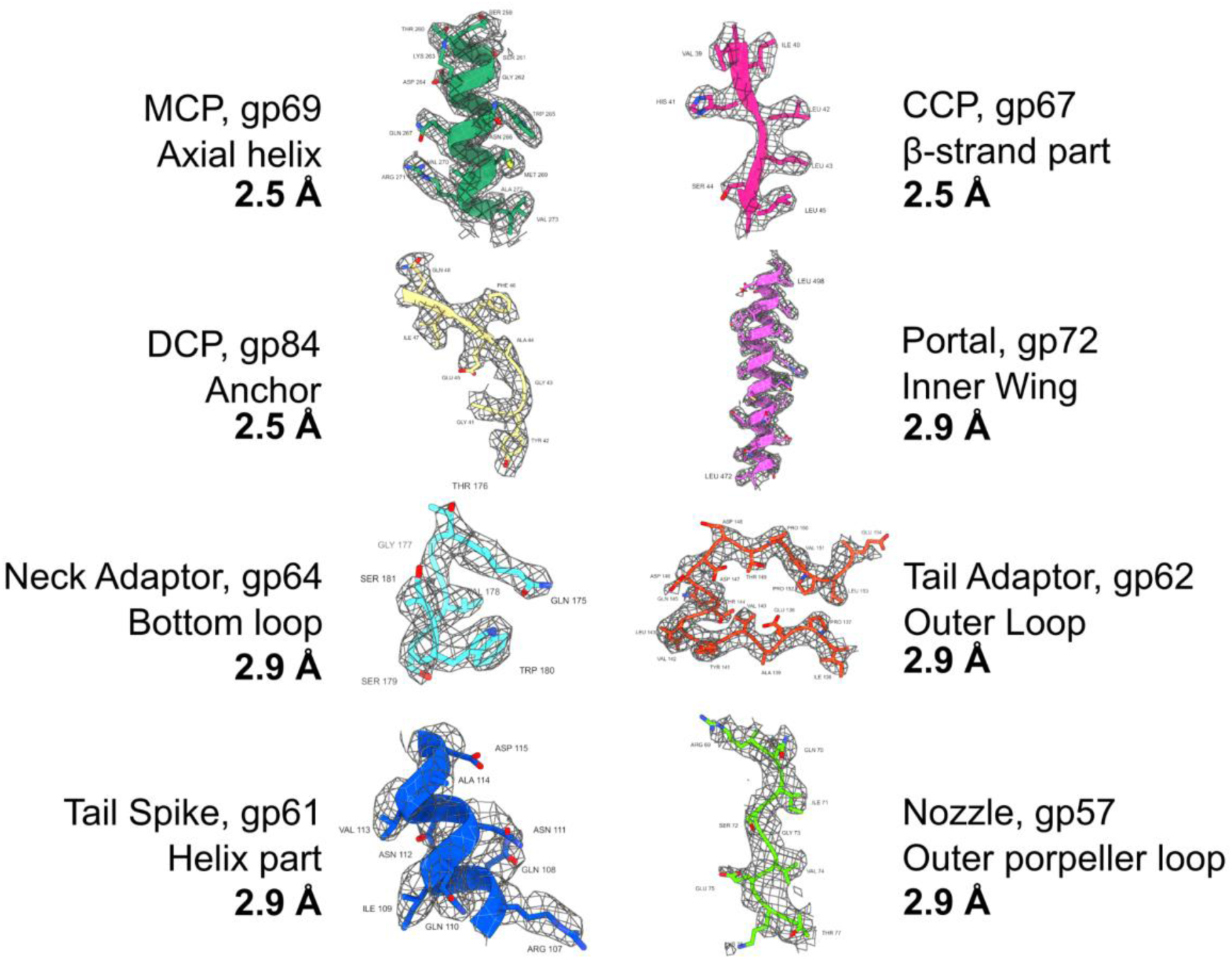
Representative fragments of cryo-EM density maps displaying the model fitting.

**Figure S7.**
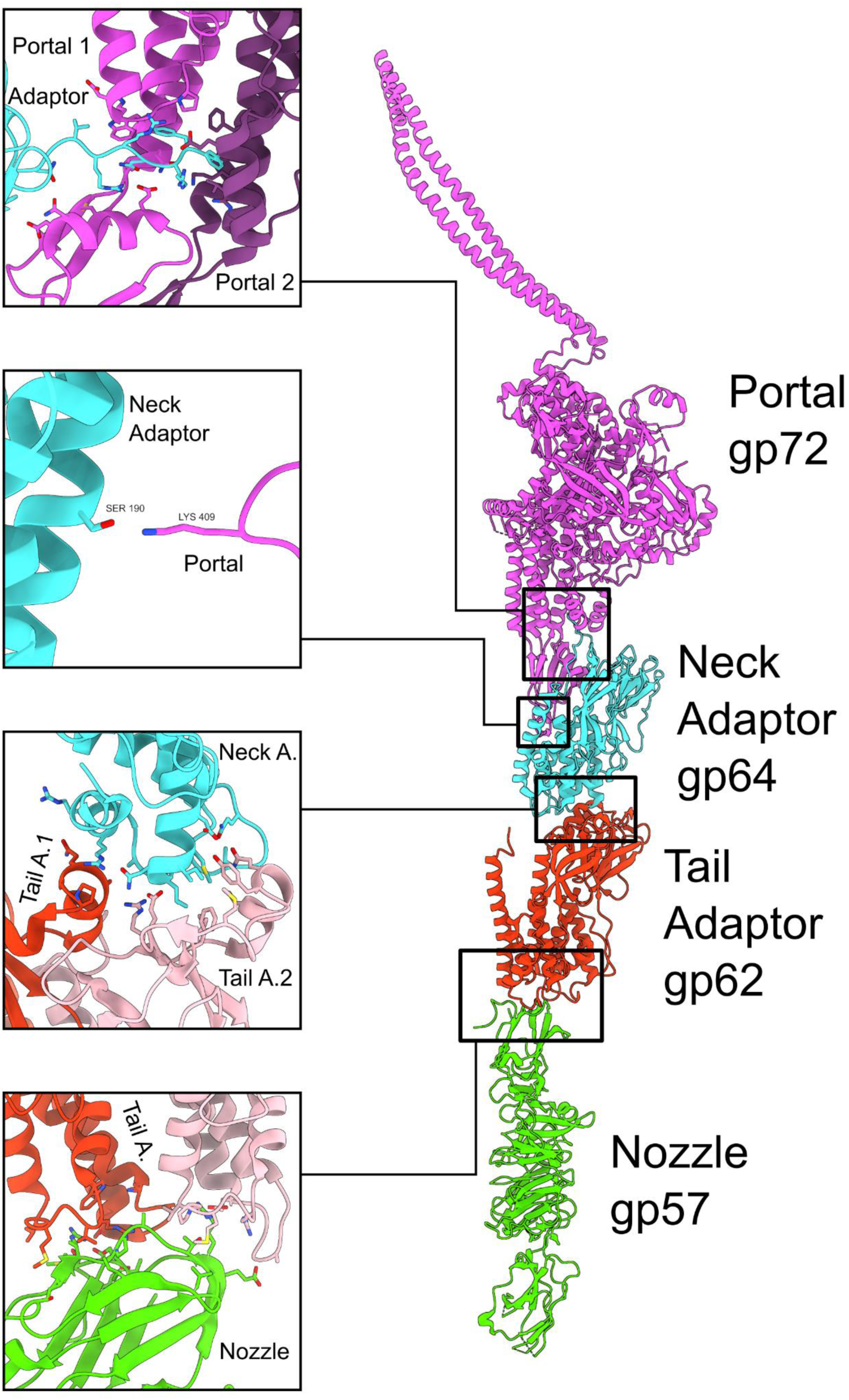
Contacts between tail tube proteins

**Figure S8.**
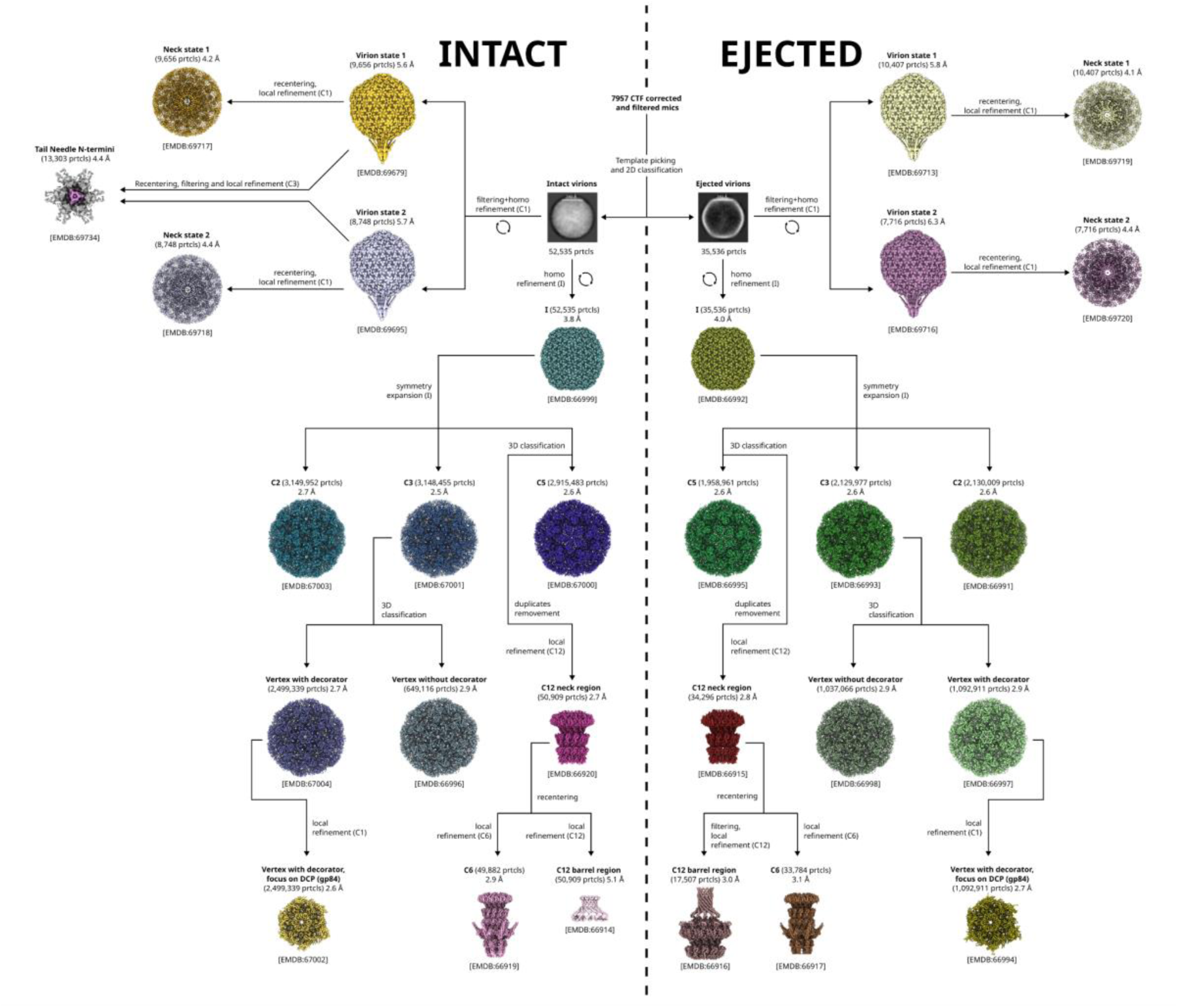
Cryo-electron microscopy single particle processing workflow.

**Figure S9.**
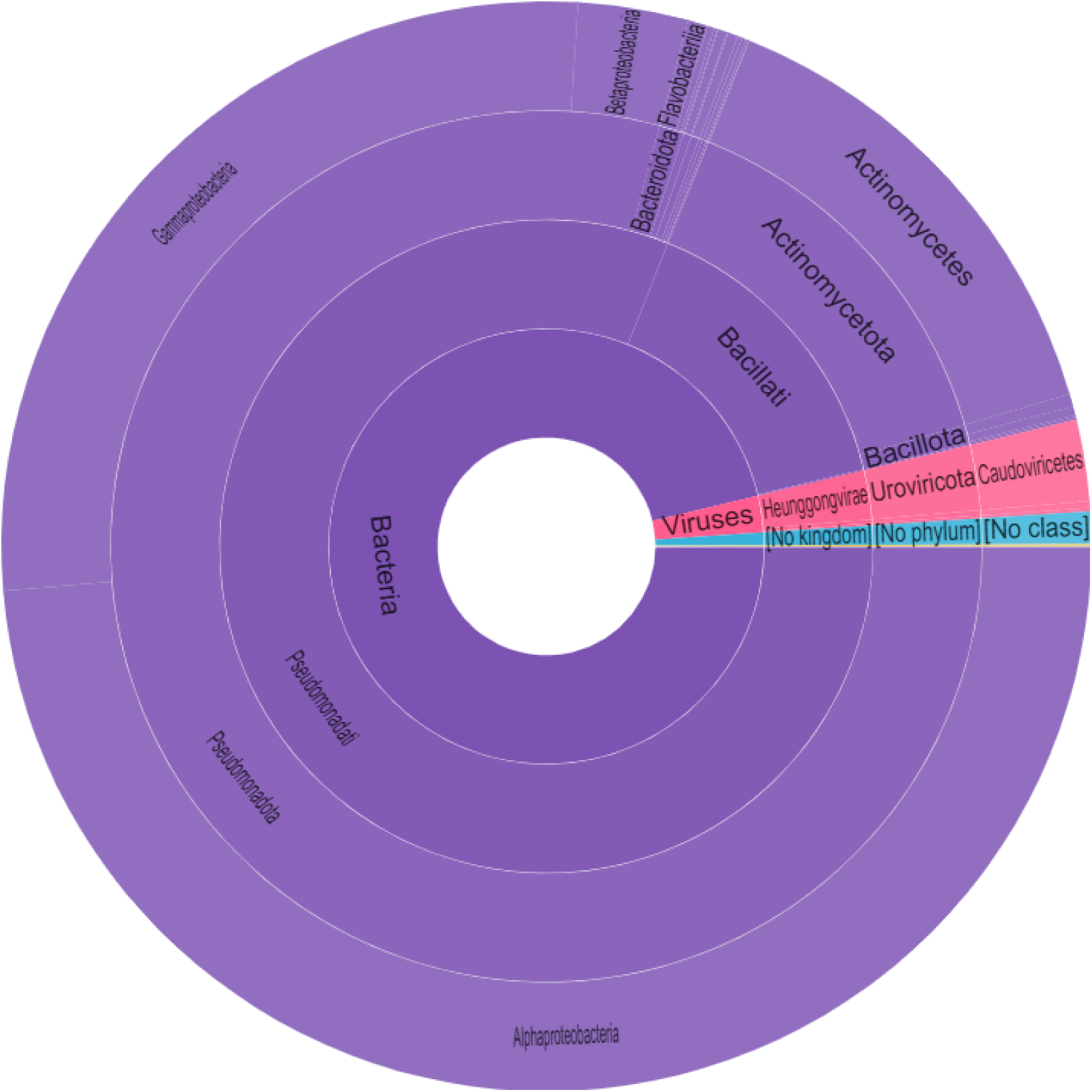
Diagram showing the distribution of DUF1737 domain in proteins of different species

